# The mitochondrial multi-omic response to exercise training across tissues

**DOI:** 10.1101/2023.01.13.523698

**Authors:** David Amar, Nicole R Gay, David Jimenez-Morales, Pierre M Jean Beltran, Megan E Ramaker, Archana Natarajan Raja, Bingqing Zhao, Yifei Sun, Shruti Marwaha, David Gaul, Steven G Hershman, Ashley Xia, Ian Lanza, Facundo M Fernandez, Stephen B. Montgomery, Andrea L. Hevener, Euan A. Ashley, Martin J Walsh, Lauren M Sparks, Charles F Burant, R Scott Rector, John Thyfault, Matthew T. Wheeler, Bret H. Goodpaster, Paul M. Coen, Simon Schenk, Sue C Bodine, Maléne E. Lindholm, the MoTrPAC Study Group

## Abstract

Mitochondria are adaptable organelles with diverse cellular functions critical to whole-body metabolic homeostasis. While chronic endurance exercise training is known to alter mitochondrial activity, these adaptations have not yet been systematically characterized. Here, the Molecular Transducers of Physical Activity Consortium (MoTrPAC) mapped the longitudinal, multi-omic changes in mitochondrial analytes across 19 tissues in male and female rats endurance trained for 1, 2, 4 or 8 weeks. Training elicited substantial changes in the adrenal gland, brown adipose, colon, heart and skeletal muscle, while we detected mild responses in the brain, lung, small intestine and testes. The colon response was characterized by non-linear dynamics that resulted in upregulation of mitochondrial function that was more prominent in females. Brown adipose and adrenal tissues were characterized by substantial downregulation of mitochondrial pathways. Training induced a previously unrecognized robust upregulation of mitochondrial protein abundance and acetylation in the liver, and a concomitant shift in lipid metabolism. The striated muscles demonstrated a highly coordinated response to increase oxidative capacity, with the majority of changes occurring in protein abundance and post-translational modifications. We identified exercise upregulated networks that are downregulated in human type 2 diabetes and liver cirrhosis. In both cases HSD17B10, a central dehydrogenase in multiple metabolic pathways and mitochondrial tRNA maturation, was the main hub. In summary, we provide a multi-omic, cross-tissue atlas of the mitochondrial response to training and identify candidates for prevention of disease-associated mitochondrial dysfunction.

## Introduction

Mitochondria are the essential powerhouses of eukaryotic cells. They generate over 90% of the ATP required for mammalian cellular homeostasis via oxidative phosphorylation. Indeed, through ATP production, their sensitivity to the cellular redox state and other signaling actions, mitochondria support and regulate many cellular processes, including steroid biosynthesis, ketone body generation, gluconeogenesis, ion homeostasis, cellular calcium signaling and programmed cell death^1^. Novel functions of mitochondrial proteins were recently discovered^2^, affirming the mitochondria as central hubs of metabolism. The circular mitochondrial DNA (mtDNA) encodes only 13 proteins, thus the majority of mitochondrial proteins are nuclear-encoded^3^.

A critical characteristic of mitochondria is their ability to adapt to subcellular-, cellular- and tissue-specific metabolic demands^4,5^. For example, mitochondria of cardiac and skeletal muscle have high capacities for aerobic ATP production to support increased energy demands during contraction/exercise^6^. In the liver, mitochondria support gluconeogenesis, ketone production and fatty acid synthesis and oxidation, and key functions that allow the liver to contribute to systemic energy substrate supply at rest, fasting or during physiological stress^7^. In brown adipose tissue, uncoupling of the inner mitochondrial membrane by uncoupling protein results in futile cycling and thermogenesis which are a critical adaptation for maintaining temperature during cold stress^8^. Given their central role in cellular homeostasis, it is perhaps not surprising that mitochondrial diseases are the most common form of inherited metabolic disorders^9^, and mitochondrial function has been linked to human health and disease risk, including type 2 diabetes, obesity, cancer, neurodegeneration, fatty liver and hepatosteatosis and cardiovascular diseases^10–13^.

Endurance exercise training is a repetitive stressor that results in robust increases in mitochondrial volume and ATP-generating capacity in the skeletal muscle. Indeed, training-induced increases in skeletal muscle mitochondrial size and numbers and improvements in substrate oxidation have been known for many decades^14,15^. Moreover, endurance exercise training imparts a myriad of health benefits through its multipotent effects across tissues and cell types, many of which are likely mediated by adaptations in mitochondrial function. Despite this, while endurance exercise training improves mitochondrial quantity and quality in skeletal muscle^16^ and liver^17^, the effects in other tissues remain largely unknown, and system-wide multiomic changes specific to the mitochondria have not been investigated.

The overall goal of the Molecular Transducers of Physical Activity Consortium (MoTrPAC) is to map the multi-omic response to exercise and training across tissues^18^. We have recently characterized the multi-omic changes across tissues in 6-month old female and male rats that were endurance exercise trained on a treadmill for 1, 2, 4 or 8 weeks^19^. Here, we focus on characterizing the longitudinal training response of mitochondrial analytes. Importantly, we included many tissues that have not been studied in detail before. Moreover, the majority of efforts in this area have included one sex only, generally studied at a single timepoint. Here, we studied male and female rats across 4 timepoints, enabling insight into time-course and sexually dimorphic mitochondrial responses to endurance exercise training. We generate a molecular map of the multi-omic mitochondrial response to endurance training across 19 different tissues and provide translational value of our maps by comparing our results to differential gene signatures of human disease, revealing gene networks that are induced by exercise but are downregulated in disease.

## Results

### Endurance training alters biomarkers of mitochondrial volume across tissues

Male and female 6-month old F344 rats were subjected to progressive endurance training on a motorized treadmill for 1, 2, 4 or 8 weeks. Additional sex-matched sedentary animals were collected as controls. Comparing trained animals to controls revealed increased aerobic capacity (VO_2_max) after 8 weeks of training in both males and females. Decreased body fat percentage was observed in males, whereas trained females retained their body fat percentage, preventing the increase that occurred over 8 weeks in the sedentary control females. The detailed training protocol and accompanying phenotypic adaptations have been described elsewhere^19^. Blood, plasma and 18 solid tissues were collected 48 hours after the last exercise bout. Samples were profiled for epigenomics using transposase-accessible chromatin using sequencing (ATAC-seq) and reduced representation bisulfite sequencing (RRBS), transcriptomics using RNA sequencing (RNA-seq), proteomics and post-translational modifications (phosphoproteome, acetylome and ubiquitylome) using LC-MS/MS, and metabolomics/lipidomics using using up to seven targeted platforms and six untargeted platforms (Fig. 1A, Methods). Transcriptomics and metabolomics/lipidomics were conducted on all 19 tissues, while proteomics and epigenomics were performed on selected tissues only (Fig. S1A)^19^. Assay-specific details, including processing pipelines, quality control, normalization and differential analyses are summarized in Methods.

**Figure 1.**
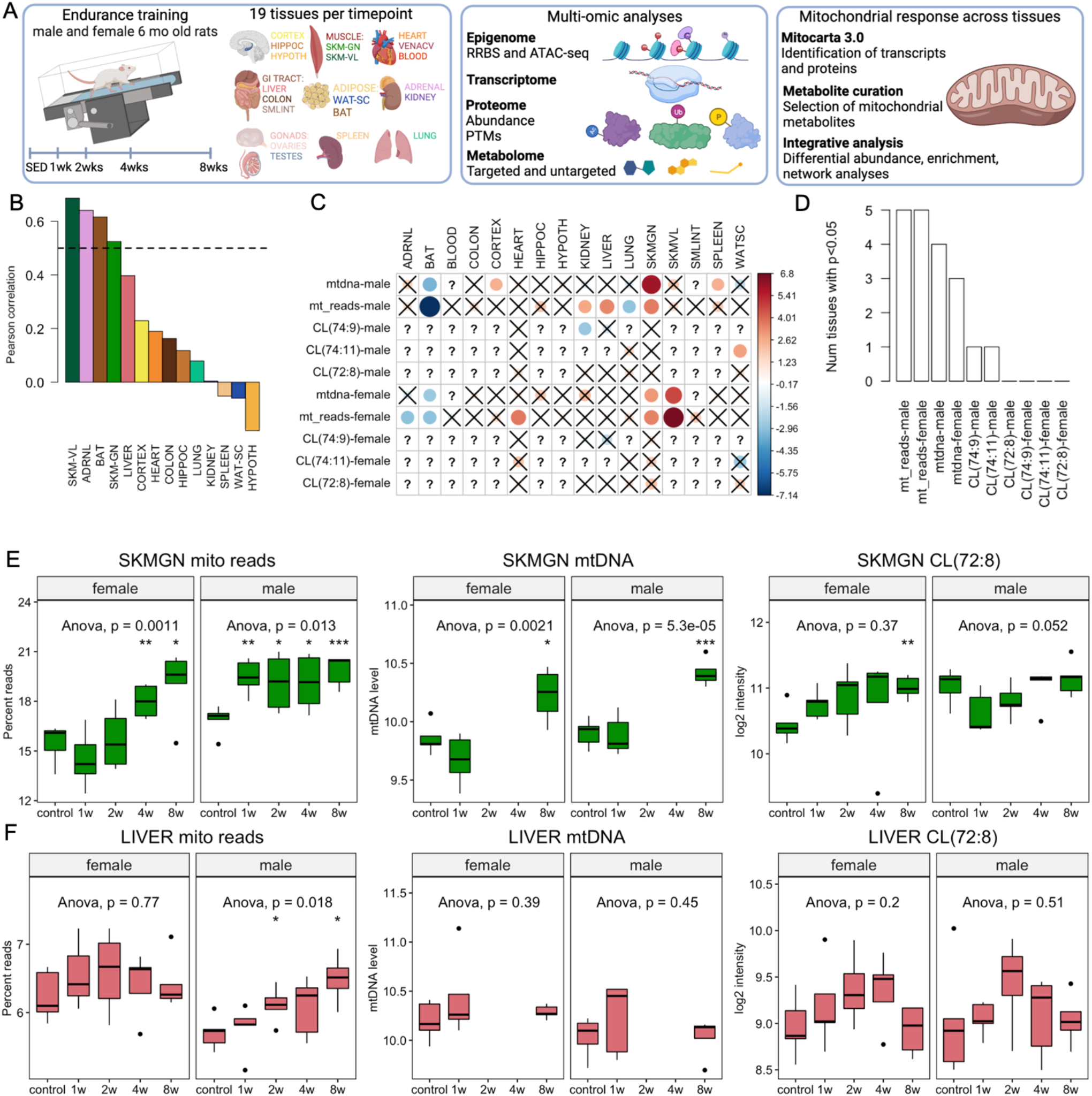
Training-induced changes in biomarkers of mitochondrial volume. **A)** Overview of the experimental design. Fischer 344 rats were subjected to a progressive treadmill training protocol. 19 tissues were collected from male and female rats that remained sedentary (SED) or completed 1, 2, 4, or 8 weeks of exercise training (tissues were harvested 48 hours after the last exercise bout). The tissues were assayed for epigenomics (8 tissues), transcriptomics (19 tissues), proteomics (7 tissues), post-translational modifications (phosphoproteome on 7 tissues and acetylome on 2 tissues), and metabolomics (19 tissues). Mitochondria-associated transcripts and proteins were selected using MitoCarta 3.0 and mitochondrial metabolites from a previously published dataset (ref. 24). HIPPOC = Hippocampus, HYPOTH = Hypothalamus, SMLINT = Small Intestine, SKM-GN = *Gastrocnemius* Skeletal Muscle, SKM-VL = *Vastus Lateralis* Skeletal Muscle, WAT-SC = Subcutaneous White Adipose Tissue, BAT = Brown Adipose Tissue, VENACV = Vena Cava. Created using BioRender.com. **B)** Correlation between mtDNA quantification and the percentage of mitochondrial RNA-seq reads across tissues. Dashed line represents rho=0.5. **C)** Training response of biomarkers of mitochondrial volume after 8 weeks of training. Cells marked with X are not significant (p>0.05). Cells marked with a ? represent tissues in which the biomarker was not assessed. Color scale is proportional to the ANOVA-test z-score. **D**) Comparison of the number of significant training responses of the mitochondrial biomarkers (p<0.05). **E-F)** Visualization of biomarker data in SKM-GN (E), and liver (F). Each boxplot represents the abundance level in a specific sex and time group. ANOVA statistics are provided for each tissue and sex combination. The whiskers extend from the hinge to the largest and lowest values, but no further than 1.5 * (the interquartile range).

To determine changes in mitochondrial volume with training, we quantified three standard biomarkers of mitochondrial quantity: (1) mtDNA copy number normalized to nuclear DNA^20^, (2) the percent of RNA-seq reads mapped to the mitochondrial genome, and (3) cardiolipin content. Cardiolipins are mitochondrial membrane-specific lipids that correlate well with mitochondrial volume^21^. Mitochondrial DNA quantification was limited to 15 tissues in animals trained for 0, 1, or 8 weeks. In contrast, the RNA-seq biomarker was available in all animals and assayed tissues; the cardiolipin data was available in six tissues in all animals: brown adipose tissue (BAT), kidney, liver, lung, *gastrocnemius* muscle (SKM-GN), and subcutaneous white adipose tissue (WAT-SC).

Striated muscle, brain and BAT were rich in mitochondria, in concordance with previous studies^22,23^. In contrast, biomarker analysis at baseline indicates a lower relative mitochondrial abundance in spleen, WAT-SC and lung (Fig. S1B-C). The percentage of mitochondrial RNA-seq reads is, by definition, a relative measurement of the abundance of the mitochondrial-encoded genes compared to all other transcripts. Thus, alternation in this relative abundance may be explained either by changes in mitochondrial volume or by independent alterations of transcriptional regulation. In our data, it was markedly correlated with mtDNA (*rho* >0.5) in four tissues: SKM-GN, *vastus lateralis* muscle (SKM-VL), adrenal and BAT (Fig. 1B). Therefore, for downstream analyses we used it as a proxy for mtDNA in these tissues. We observed that different cardiolipins tended to be highly correlated across the samples from the same tissue (Fig. S1D-F).

We observed a significant response to training in at least one of the mitochondrial biomarkers in 15 of the 19 tissues (Supplementary data S1). Focusing on week 8, the percent of mitochondrial reads responded in eight tissues, with a concordant direction of response with mtDNA (Fig. 1C-D). Moreover, the top responding sex-tissue combinations manifested a concordant response of the mitochondrial-encoded transcripts (Fig S2), suggesting potential changes in both mtDNA quantity and transcriptional activity. Specifically, global upregulation of these mitochondrial-encoded genes was consistent with upregulation in both males and females for all three biomarkers in skeletal muscle (Fig. 1E, S2L). In contrast, the liver manifested a mild response that was limited to upregulation in males (Fig. 1F, see Fig. S3 and Supplementary data S1 for all biomarker data). In summary, we describe substantial cross-tissue differential changes of mitochondrial biomarkers in response to training, providing estimation of change in mitochondrial volume that were used for additional analyses.

### The mitochondrial multi-omic response to exercise training is tissue-specific

We utilized MitoCarta 3.0^3^ to identify mitochondria-associated genes and proteins, and data from Heden et al.^24^ to select mitochondrial metabolites. Principal component analysis (PCA) of the baseline mitochondria-associated transcriptome showed a clear primary separation by tissue, with additional separation by sex for multiple tissues (Fig. S4A). The baseline mitochondrial proteome was more similar across analyzed tissues, with distinct sex differences observed in WAT-SC and SKM-GN (Fig. S4B).

Training regulated mitochondria-associated analytes across all -omes and tissues. In total, 719 genes, corresponding to 63% of all mitochondria-associated transcripts, and 513 proteins (38% of all mitochondria-associated proteins) significantly changed with training in at least one sex and tissue. The most responsive tissues (>10% differential transcripts and/or proteins) were the adrenal gland, BAT, blood, colon, heart, liver, SKM-GN, SKM-VL and WAT (Fig. 2A). SKM-GN, liver and heart showed the greatest proteomic response, while the metabolome/lipidome changed the most in blood (plasma), heart and liver. In contrast, there was mild mitochondrial response in the three investigated brain regions (cortex, hippocampus and hypothalamus), kidney, lung, testes and vena cava (Fig. 2A). Interestingly, the lung had widespread changes in non-mitochondrial analytes^19^. The lungs and vena cava are tissues that experience large increases in blood flow with exercise but little change in metabolic demand. While there were no significant pathways enriched in mitochondria-associated analytes in the lung, we observed a decreased expression of superoxide dismutase (*Sod2*), similar to a previous study in rats on isolated lung mitochondria^25^. Very few significant endurance training related changes in protein phosphorylation and ubiquitination of mitochondrial proteins were observed across tissues. Comparing the differential transcripts of the top nine responsive tissues demonstrated a largely tissue-specific response, with moderate overlap that was nevertheless significant for multiple pathways (Fig. 2B). Similar moderate-low overlap was also observed at the protein level (Fig S4C). The greatest transcriptional response was observed in the adrenal gland, followed by BAT and colon, where the commonly regulated genes (m=28) encoded tricarboxylic acid cycle (TCA) enzymes and subunits of Complex I (Fig. 2B).

**Figure 2.**
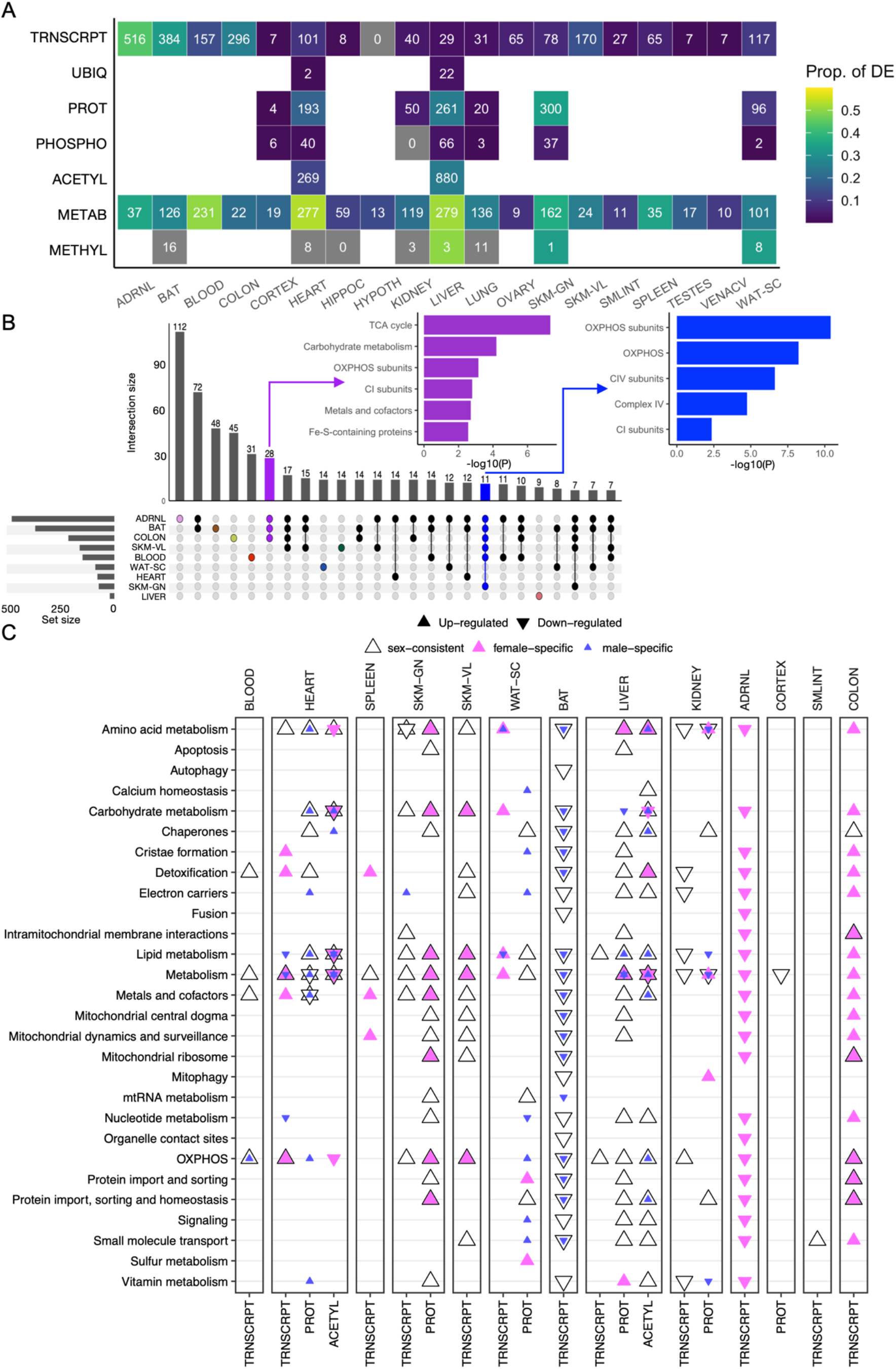
The multi-omic mitochondrial response to training across tissues. **A)** Heatmap of the number of mitochondria-associated analytes that significantly changed in abundance over the training time course in at least one sex (5% FDR). Each cell represents results for a single tissue and data type. Numbers indicate the number of training-differential mitochondrial analytes and colors indicate the proportion of measured analytes identified in MitoCarta that are differential. **B)** UpSet plot of the training-differential MitoCarta transcripts across tissues. Numbers above vertical bars indicate the number of transcripts differentially regulated by training in the tissues indicated by connected points below the bar. Horizontal bars indicate the total number of differential transcripts in each tissue. Pathway enrichment results using the MitoCarta pathways are shown for the colored bars; purple represents the 28 differential genes that were common in the adrenal glands, BAT and colon, whereas the blue bar represents 11 differential genes that were common among six tissues. **C)** MitoCarta pathway enrichments for the 8-week training timepoint in the 9 tissues that showed the greatest mitochondrial training response. The 8-week male and female differential transcripts were identified using our graphical analysis (see Methods). The plot shows the top pathway from each MitoCarta subcategory with the greatest number of enrichments. This was determined by taking the sum-of-log combined p-value per tissue and pathway. Each point represents a significant pathway enrichment in a given node, where the direction of the triangle indicates the direction of the training effect (up or down) and the color indicates the corresponding sex (blue = male, pink = female, black triangle indicates sex-consistent enrichment).

The differential analysis above mainly focused on an overall training response across sex and time. Subsequently, we used the same models for contrasting each training time point with its sex-matched controls. This analysis produced time- and sex-specific summary statistics including fold changes and z/t-scores (see Methods). We also utilized a graphical approach for representing these differential abundance analysis results in all tissues. This analysis takes these timewise z-scores from males and females, and uses the *repfdr* empirical Bayes algorithm^26^ to identify the main trajectories in a selected set of tissues and omes. These are used for visualization, identifying sets of analytes that change in a specific time- and sex combination, and mitochondria-specific pathway enrichment of these sets^19^ (see Methods).

Using the graphical analysis results, we first focused on the long-term adaptation in week 8. MitoCarta^3^ pathway enrichment analysis identified a consistent multi-omic upregulation of numerous mitochondrial pathways in the heart, skeletal muscles, WAT-SC and liver (Fig. 2C). In the striated muscle tissues, the majority of the mitochondrial adaptation was sex-consistent. The liver response was also similar between the sexes, specifically in the proteome and acetylome, with very few transcriptional changes. Many mitochondrial pathways also increased in trained WAT-SC, but showed mostly sex-specific regulation. Training specifically induced carbohydrate and sulfur metabolism pathways in female WAT-SC, whereas induction of oxidative phosphorylation (OXPHOS) and proteins associated with cristae formation and calcium homeostasis were specific to males.

Epidemiological evidence of reduced risk for colon cancer in trained individuals^27^ supports an effect of training on the colon. Interestingly, we observed a dynamic mitochondrial response, with an overall downregulation of mitochondria-associated transcripts after the first two weeks of training, and a delayed upregulation after 8 weeks (Fig. S5A-B). Importantly, many exercise interventions in animal studies are terminated at 4 weeks, where virtually no transcriptional changes were observed in our data. The overall response was substantially greater in females compared to males. Pathway enrichment analysis of the female-specific transcript changes showed induction of oxidative phosphorylation (all electron transport chain/ETC complexes apart from Complex II) and TCA cycle genes in response to 8 weeks of training (Fig. S5C).

However, despite observing a greater differential response in females, these transcripts converged to similar values as males by week 8 (i.e., these had lower expression at baseline, see Fig. S5D). The majority of the mitochondrial proteome is nuclear-encoded, and import of presequence proteins through the mitochondrial membranes is therefore a critical function regulating mitochondrial biogenesis and homeostasis^28^. Sex-consistent upregulation was observed for the presequence import TIM23, where *Timm23* and *Timm50* were significant at the gene level. TIM23 is stabilized by cardiolipins in the mitochondrial membrane, thus the combined increase in TIM23 and cardiolipin synthesis genes (*i*.*e. Agpat4, Prelid1* and *Ptpmt1*) after 8 weeks of training suggests an altered mitochondrial membrane dynamics in the colon. Thus, we observe sexual dimorphism in the mitochondrial endurance training responses in the adrenal, colon and white adipose tissues, and a largely tissue-specific mitochondrial training response overall.

### Changes in mitochondrial volume partially explain the training response in the adrenal gland and brown adipose tissue

Differential abundance of mitochondrial analytes can be explained by a change in mitochondrial composition/function, volume or by a combination of both. To determine which of the differential analysis results can be explained by changes in mitochondrial volume we repeated the differential analysis in eight tissues with adjustment for: (1) cardiolipin measured in six tissues (heart, kidney, liver, lung, SKM-GN, and WAT-SC), and (2) percent of mitochondrial RNA-seq reads (an mtDNA proxy in adrenal, BAT, SKM-GN, and SKM-VL). Overall, adjustment primarily affected transcript changes, whereas proteomic results had substantially fewer modifications to the sets of selected analytes (Fig. S6A).

We next compared the timewise-specific results. For each analyte we used the pre- and post-adjustment models and extracted the timewise z-scores, comparing each time point to its sex-matched controls. This resulted in 16 z-scores: four time points before and after adjustment for each sex. We observed that most differential analysis results did not change substantially after adjustment, with only 799 analytes out of the 2,167 analyzed analytes (37%) producing a difference of three or greater in at least one of the time points (Extended Data S2). Clustering analysis of these 799 z-score trajectories identified four clusters using the k-means algorithm (Supplementary data S2). Fig. 3A shows the cluster trajectories, and Fig. 3B shows the cluster composition by tissue and -ome. Cluster 1 represents a downregulation pattern that is specific to BAT, whereas clusters 2 and 4 represent two responses of the adrenal gland. Cluster 3 was dominated by SKM-VL, demonstrating male differential transcripts at 8 weeks that were largely driven by changes in mitochondrial volume. The few genes that were explained by changes in mitochondrial volume in SKM-GN were mitochondria-encoded (Supplementary data S2). Differential genes affected by adjustment in SKM-VL were mainly encoding ETC components and a few TCA components, while genes associated with mitochondrial dynamics and amino acid metabolism were robust to adjustment for mitochondrial volume.

**Figure 3.**
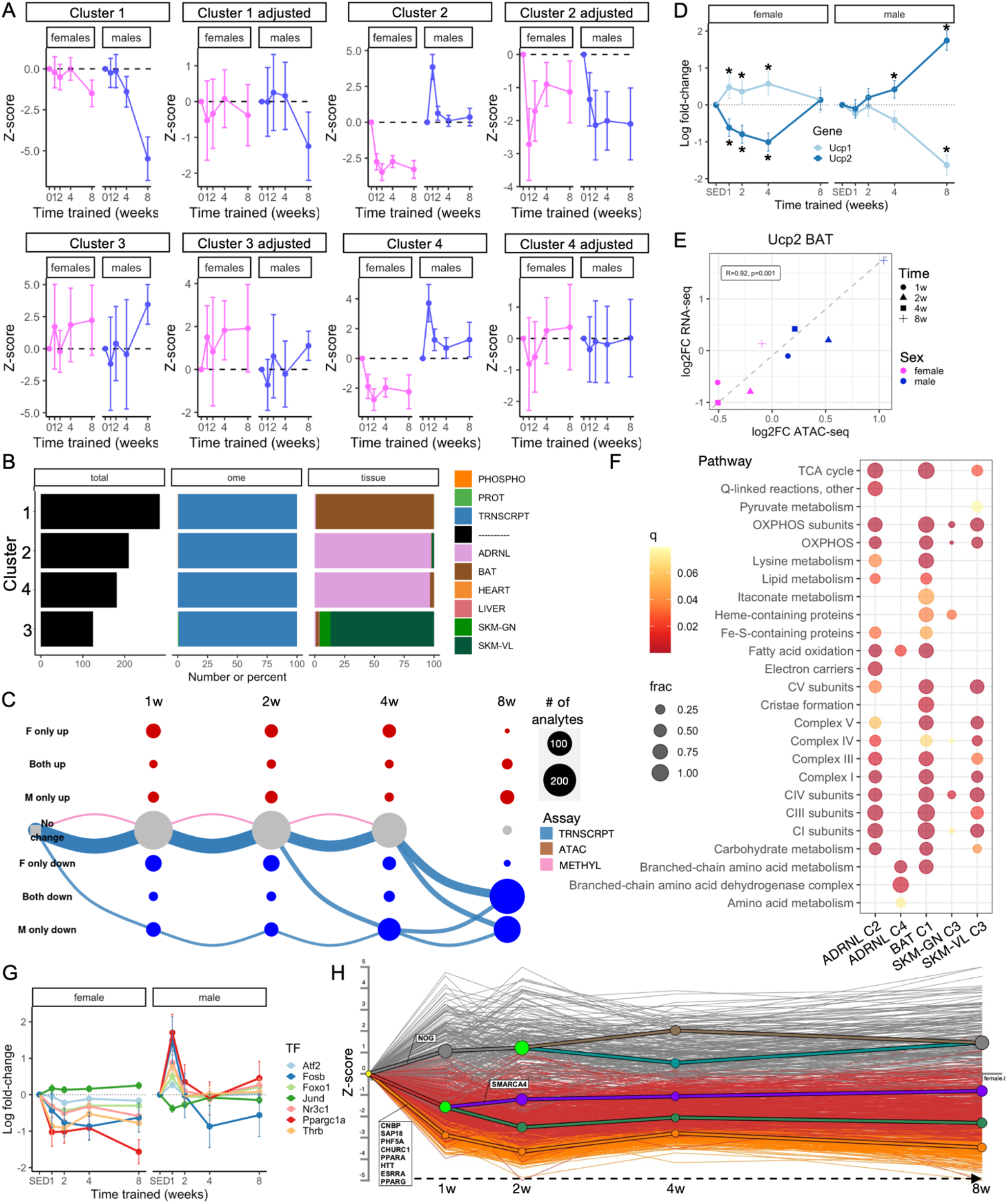
Changes in mitochondrial volume partially explain the training response in the adrenal gland and brown adipose tissue. **A)** Clustering analysis showing patterns affected by adjustment for biomarkers of mitochondrial volume. **B)** Characteristics of the clusters in A. Number of total analytes, distribution across –omes and tissues is illustrated for each cluster. **C)** Graphical representation of the mitochondria-associated training-differential analytes in brown adipose tissue (BAT). Each node represents one of nine possible states (row labels, with F for females and M for males, seven states shown) at each of the four sampled training time points (column labels). Edges are drawn between nodes to represent the path of differential analytes over the training time course, with color representing the –ome. This graph includes the five largest paths for the BAT. Both node and edge size are proportional to the number of analytes represented by the node or edge. **D)** Gene expression changes (log2 fold-change) of Ucp1 and Ucp2 in females (left) and males (right). * indicates significant timewise change (FDR<0.05). **E)** Correlation between changes (log2 fold-change) in Ucp2 expression and chromatin accessibility (intronic region of UCP2, chr1:165508254-165509507). Each point represents the average for n=5 animals assayed for that timepoint and sex. **F)** Pathway enrichment analysis of the clusters affected by mitochondrial volume from A using the MitoCarta 3.0 Pathways. Color indicates significance of the enrichment (q-value) and size indicates fraction of analytes in the cluster that is part of the pathway. **G)** Gene expression changes (log2 fold-change) of examples of known PPARGC1A interactors. All are significantly upregulated in males after 1 week and downregulated in females after 8 weeks, with exception of Jund, which is significantly regulated in the opposite directions (FDR<0.05). **H)** Dynamic regulatory events miner (DREM) analysis results of the MitoCarta genes in female adrenal gland predict several transcription factors to be involved in the early (1w) responses.

Cluster 1 almost exclusively comprises brown adipose transcripts, illustrating a marked decrease in males after 8 weeks (Fig. 3A), which is largely attenuated after the adjustment. Nevertheless, a slight downregulation still remains, suggesting that while some of the substantial transcriptional downregulation is due to a decrease in mitochondrial volume, several pathways may exhibit an additional relative downregulation among the mitochondrial analytes. While several studies demonstrate improved mitochondrial function with endurance training in WAT-SC, there are conflicting results in BAT^29^, with the majority of prior studies showing either no effect or a decrease in BAT mass and activity with training^30–33^. BAT is unique in its response among our investigated tissues in its overall training-induced downregulation of mitochondrial analytes (Fig. 3C). This late (week 8) downregulation is observed in major mitochondrial metabolic pathways: the TCA cycle and oxidative phosphorylation (Fig. 2C, S6B-C), suggesting a reduced metabolic capacity in BAT with endurance training. *Ucp1*, which is considered the key thermogenic uncoupler, is downregulated specifically in males and mostly in week 8, whereas the chromatin accessibility and associated gene expression of *Ucp2* are inversely correlated with *Ucp1* (Fig. 3D-E). *Ucp2* does not have the uncoupling potential of *Ucp1* but has been shown to limit glucose oxidation and instead enhance glutaminolysis^34^. Together, these results suggest a potential mechanism for energy preservation in response to training.

We used DREM (Dynamic Regulatory Events Miner)^35^ to identify potential upstream regulators of the mitochondrial response in BAT. DREM was specifically developed for time course data by combining clustering analysis with transcription factor prediction that may explain the identified trajectories. Known regulators of mitochondrial metabolic genes, including PPARA, PPARD and their coregulator PPARGC1A, are associated with the late onset of downregulation of mitochondria-associated genes in male BAT (Fig. S6D). Several predicted upstream regulators (*e*.*g*. CNBP, SAP18, CHURC1 and PHF5A) were significant also in colon and the adrenal glands, which together with BAT show the largest transcriptional changes with training.

The adrenal gland, a previously unexplored tissue for training adaptation, had the largest mitochondrial response to endurance training, with differential expression of almost 50% of all mitochondria-associated genes (Fig. 2A, S7A). In our clustering analysis, the adrenal responses were separated into two clusters that are both largely explained by changes in mitochondrial volume. In both cases, the pre-adjustment trajectories are similar, showing a consistent downregulation in females across the time points, with a marked upregulation in males in week 1 (Fig. 3A, S7B). This female-specific response covers the majority of genes involved in beta-oxidation, the TCA cycle and the ETC (Fig. S7C-D). Post-adjustment, cluster 2 manifests downregulated patterns, with more similar male and female trajectories, while cluster 4 shows a nullified effect, with most post-adjustment cluster members having z-scores close to zero (Fig. 3A, Supplementary data S2, Extended Data S2). Interestingly, these two clusters differ in their enrichment results, where the changes that are driven by a reduction in mitochondrial volume in females (cluster 4) are mainly associated with amino acid metabolism (Fig. 3F). The female-specific decreases in fatty acid metabolism, TCA cycle and ETC are largely robust to adjustment, indicating a reduced mitochondrial metabolic capacity of the female adrenal gland with training on top of a reduction in mitochondrial volume. In light of extensive mitochondrial remodeling of the adrenal gland, we interrogated the potential drivers underlying these responses. PGC1a is a key regulator of mitochondrial biogenesis, and because the main differential trajectory includes many of the known PGC1a coactivator response genes^36^, this is a likely underpinning of the observed changes. PGC1a followed the same main trajectory, increasing in males at week 1 with a decrease in females at week 8. Interestingly, several of the known interactors of PGC1a showed the same pattern, including the PPARs^19^, Fosb, Foxo1 and Thrb (Fig. 3G). Glucocorticoids, produced by the adrenal cortex, are known activators of PGC1a expression^37^, as are changes in energy balance, temperature and calcium concentration^36^. DREM predicted several transcription factors, including CNBP, SAP18, PHF5A and CHURC1, as upstream regulators in the adrenal in both sexes (Fig. 3H, S7F). In females, additional factors were predicted, including PPARA, PPARG and ESRRA (Fig. 3H), which are all interactors of PGC1a. Thus, the DREM results collectively suggest potential upstream regulators of PGC1a and mitochondrial biogenesis in the adrenal gland in response to endurance training.

In summary, we show that the sex-specific mitochondrial responses in BAT and adrenal glands are largely driven by changes in mitochondrial volume. The training-induced responses in other tissues are largely independent of changes in mitochondrial volume, thus reflecting adaptation in functional efficiency of differential pathways.

### Training induces sex-consistent mitochondrial upregulation and increased fatty acid metabolism in female skeletal muscle

The SKM-GN analytes showed sex-consistent trajectories, with an overall upregulation across the 8 weeks of training (Fig. 4A). Interestingly, most of the mitochondrial changes occured already after a single week and remain differential across the entire 8-week training period. The main trajectory was driven by alterations at the protein level, with changes in OXPHOS seen for both transcripts and proteins. In contrast, lipid metabolism and TCA cycle enzymes were almost exclusively regulated at the protein level. These findings were concordant with pathway enrichment analyses of the sex-consistent upregulated genes in both SKM-GN and SKM-VL (Fig. 4B, Fig. S8A-C). Moreover, these enrichment analyses revealed female-specific metabolic effects of training. Females specifically increased lipid metabolism transcripts in SKM-VL and fatty acid oxidation proteins in SKM-GN, e.g. ACSS1, ECHS1, ECI1 and HADHA (Fig. 4C). Similarly, the complex III pathway was only upregulated in females (Fig. S8D).

**Figure 4.**
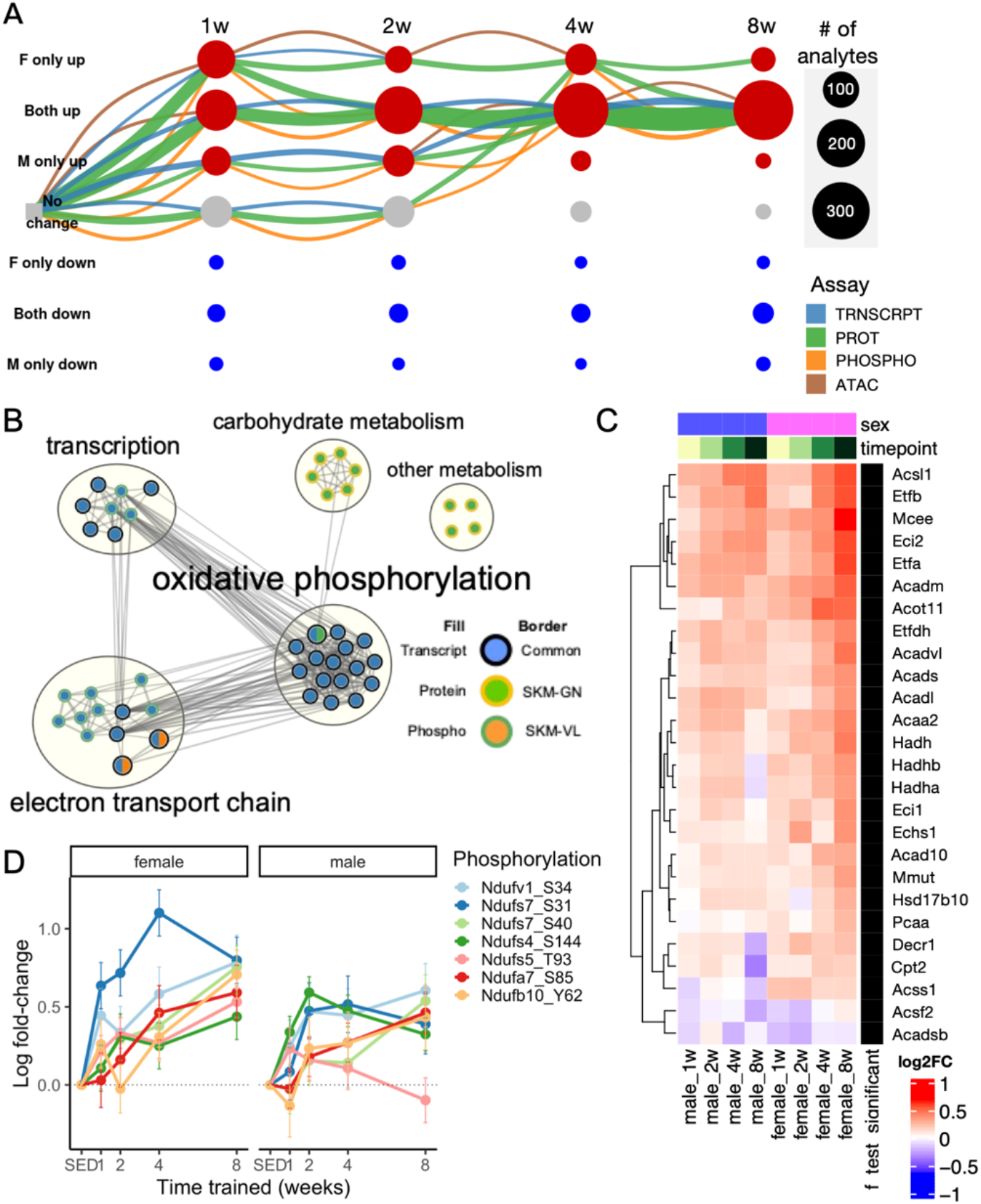
Endurance training induces largely sex-consistent increases in metabolic protein abundance in skeletal muscle. **A)** The dynamics of the molecular training response visualized by constructing a summary graph in which rows represent nine combined states (row labels, with F for females and M for males, seven states shown) and columns represent the four training time points. Nodes correspond to a combination of time, sex, and state. An edge connects two nodes from adjacent time points, representing a local temporal pattern, with edge color representing the –ome. The differential abundance trajectory of any given training-regulated analyte is represented by drawing a path through the nodes in this graph. This graph represents the mitochondria-associated training-differential analytes in the gastrocnemius (SKM-GN). This graph includes the five largest trajectories (by number of analytes). Both node and edge size are proportional to the number of analytes represented by the node or edge. **B)** Network view of pathway enrichment results corresponding to the analytes of the week 8, sex-consistent upregulation nodes in SKM-GN (A) and SKM-VL. Nodes indicate significantly enriched pathways (10% FDR), and an edge represents a pair of nodes with a similarity score of at least 0.3 between the gene sets driving each pathway enrichment. Node fill color indicates for which –ome or –omes a pathway is significant, while border color indicates if the pathway is significant in one or both skeletal muscle tissues. Node size is proportional to the number of differential analyte sets (e.g., vastus lateralis transcripts) for which the pathway is significantly enriched. Clusters of enriched pathways were defined using Louvain community detection, and are annotated with high-level biological themes. **C)** Fatty acid oxidation pathway enrichment for the gastrocnemius (SKM-GN) proteome. Only significant genes are shown. Rows are clustered using hierarchical clustering. **D)** Log2 fold changes of significant differential protein phosphorylation sites in Complex I proteins in males and females. All phosphorylation changes are significant in females, whereas all except Ndufs5_T93 are significant in males after 8 weeks of training.

Phosphoproteome changes have been described in skeletal muscle in response to acute exercise^38^ but are lacking for long-term training. A large percentage of Complex I proteins are regulated by phosphorylation^39,40^, and increased phosphorylation of Complex I has been associated with increased activity in cardiac muscle^41^, although site-specific effects on individual enzyme activities are unknown. We observed an induction of Complex I phosphorylation in skeletal muscle with training (retained 48 hours after the last bout of exercise) (Fig. 4D), suggesting phosphorylation as a regulatory mechanism for increased ETC activity in response to long-term training. The abundance of several of the Complex I proteins also changed with training, but to a lesser extent compared to phosphorylation. Thus, we observe several levels of regulation associated with increasing oxidative capacity in skeletal muscle, which is critical for preventing age-related decline in mitochondrial function^42^.

### Endurance training alters the cardiac mitochondrial acetylome

Differential analysis of cardiac mitochondrial analytes revealed a sex-consistent upregulated trajectory of transcripts and proteins across 8 weeks of training (Fig. 5A). The second largest trajectory showed delayed downregulation at week 8 (Fig. 5A-B). Training increased transcript abundance of OXPHOS genes, similar to skeletal muscle, while TCA, BCAA (branched chain amino acid) and fatty acid metabolism pathways were regulated across multiple -omes (Fig. 5C, and Fig. S9A-B). These findings were corroborated by significant enrichment of cardiac tissue TCA acids and amino acids. Sex-consistent downregulated proteins were associated with Coenzyme Q metabolism, whereas a reduction in acetylation occurred for BCAA and lipid metabolism proteins (Fig. 5D).

**Figure 5.**
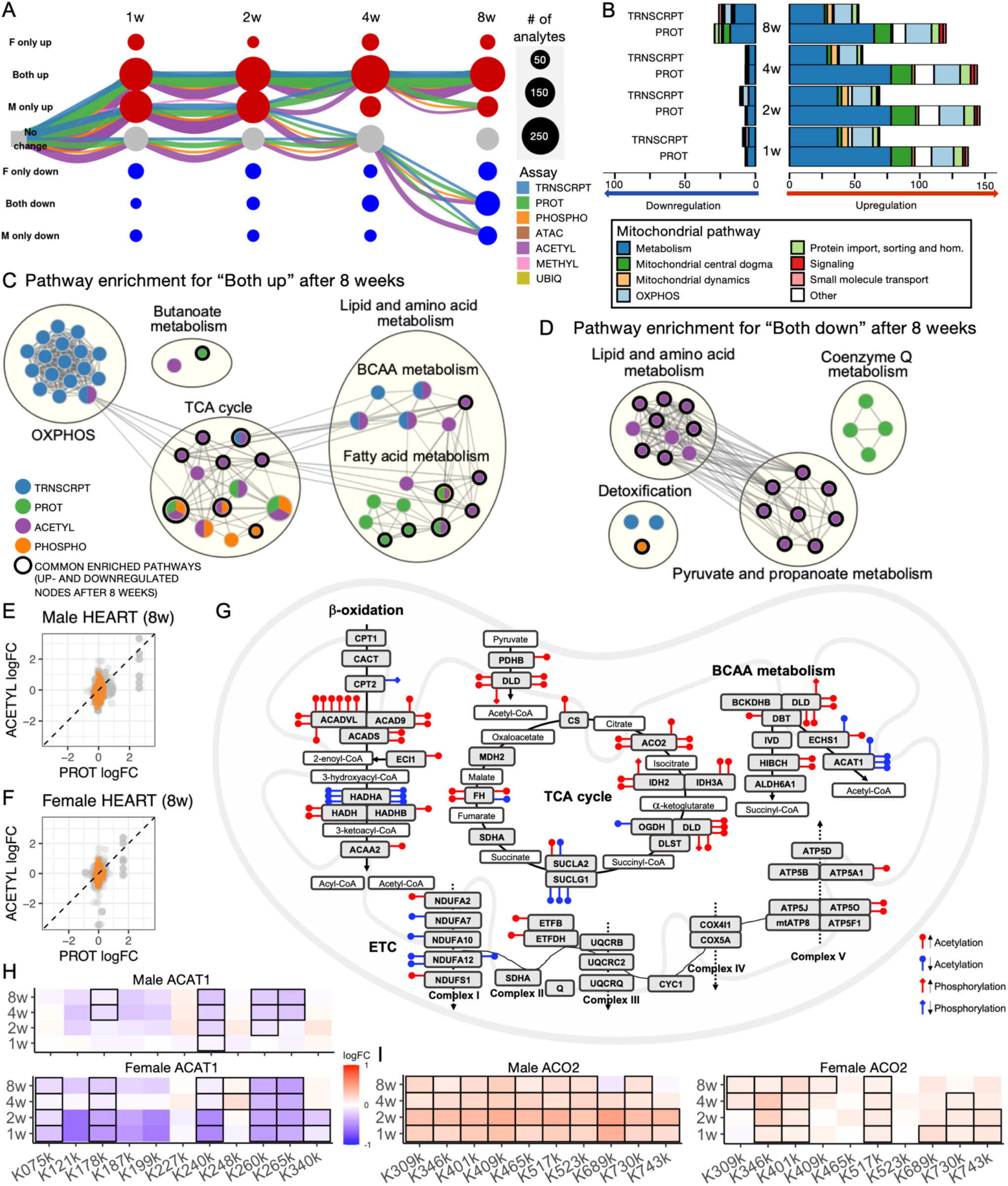
Endurance training alters the cardiac mitochondrial acetylome. **A)** Graphical representation of the mitochondria-associated training-differential analytes in the cardiac muscle. Each node represents one of nine possible states (row labels, with F for females and M for males, seven states shown) at each of the four sampled training time points (column labels). Edges are drawn between nodes to represent the path of differential analytes over the training time course, with color representing the –ome. This graph includes the five largest paths for cardiac muscle. Both node and edge size are proportional to the number of analytes represented by the node or edge. **B)** Number of significantly up- and downregulated mitochondria-associated cardiac transcripts and proteins at each training timepoint, with color representation based on the main MitoCarta pathway association of each analyte. **C-D)** Network view of pathway enrichment results corresponding to the analytes **C)** downregulated in both sexes after 8 weeks (the 8w_F-1_M-1 node in (A)) and **D)** upregulated in both sexes after 8 weeks (the 8w_F1_M1 node in (A)). Nodes indicate significantly enriched pathways (10% FDR), and an edge represents a pair of nodes with a similarity score of at least 0.3 between the gene sets driving each pathway enrichment. Node fill color indicates for which –ome or –omes a pathway is significant, while a black border color indicates if the pathway is significant in both the down- and upregulated nodes. Node size is proportional to the number of differential analyte sets for which the pathway is significantly enriched. Clusters of enriched pathways were defined using Louvain community detection, and are annotated with high-level biological themes. **E-F)** Correlation between changes (log2 fold change) in protein levels and acetylation levels in males (E) and females (F). Orange color indicates MitoCarta proteins, while other proteins are shown in grey. **G)** Significant acetylation and phosphorylation changes (FDR<0.05) of mitochondrial metabolic proteins in male and female cardiac muscle after 8 weeks of endurance training (sites changing in only one sex are not illustrated). Each lollipop represents a specific acetylation (rounded top) or phosphorylation (diamond top) site, where red color indicates increases and blue decreases. Multiple lollipops on the same protein indicates several sites significantly changed with training. Hadha had 8 differentially acetylated sites in total, out of which only 6 are illustrated due to space constraints. Proteins displayed with a name different than the official gene name are Atp5b = Atp5f1b, Atp5c = Atp5f1c, Atp5a1 = Atp5f1a, Atp5j = Atp5pf, Atp5f1 = Atp5pb, Atp5o = Atp5po. **H)** Site-specific acetylation changes in ACAT1 in males (top panel) and females (bottom panel), and in **I)** ACO2 in males (left panel) and females (right panel). All displayed sites were differentially acetylated overall (taking all timepoints and sexes into account, FDR<0.05), and sites that reach timewise significance (FDR<0.05) are highlighted with black frames.

Endurance training induced multiple acetylation changes of mitochondrial proteins (220 significant sites after 8 weeks of training, Supplementary data S3); a known regulatory mechanism of cardiac bioenergetics that has been associated with cardiac disease pathogenicity^43^ and cardiac aging^44^. The majority of acetylation changes occurred independently of changes in protein abundance following training (Fig. 5E-F). Sex-consistent protein-specific acetylation changes in response to 8 weeks of training occurred in key bioenergetic pathways (Fig. 5G). Multiple β-oxidation, BCAA and TCA cycle enzymes were significantly regulated at multiple individual acetylation sites. Reduced acetylation was primarily observed in HADHA (Fig. S9C), NDUFA12 and ACAT1. Reduced acetylation of ACAT1, an enzyme involved in both β-oxidation and BCAA catabolism, occurred at lysines 260 and 265 (Fig. 5H), two sites where deacetylation is known to increase protein activity due to increased affinity for Coenzyme A^45^. Conversely, training induced acetylation of multiple lysines in cardiac aconitase (ACO2, Fig. 5I), which increases the activity of the TCA cycle enzyme^46^. SIRT3, the main protein deacetylase in cardiac mitochondria, showed a small but significant increase in protein expression after 8 weeks of training. Thus, our study demonstrates major acetylation changes at specific cardiac mitochondrial proteins in response to training, suggesting a novel molecular mechanism for the cardioprotective effects of exercise.

### Remodeling of the metabolic protein acetylome drives training-induced mitochondrial adaptation in the liver

Endurance training leads to functional improvements in hepatic mitochondria^47^, but the molecular mechanisms driving these changes are unknown. Eight weeks of training altered mitochondrial analytes across all -omes, with the majority of changes occurring at the level of protein and protein modifications. Very few transcriptional changes occurred (Fig. 6A), an effect that we observed in our previous MoTrPAC analysis^19^, which could be tied to higher rates of mRNA turnover in the liver^48^. The main trajectory in the liver showed male-specific increases across the first 4 weeks of training and an increase in both sexes after 8 weeks (Fig. 6A). The pathway enrichment for this trajectory was dominated by changes in acetylation of Complex V proteins, mitochondrial biogenesis and BCAA catabolism, as well as lipid metabolism proteins (Fig. S10A). The second largest trajectory, consisting of >200 analytes, remained unchanged during the first 4 weeks of training but subsequently increased in both sexes by week 8 (Fig. 6A). Here, pathway enrichment was dominated by changes in mitochondria-encoded transcripts of Complex I (Mt-Nd1-6) (Fig. S10B).

**Figure 6.**
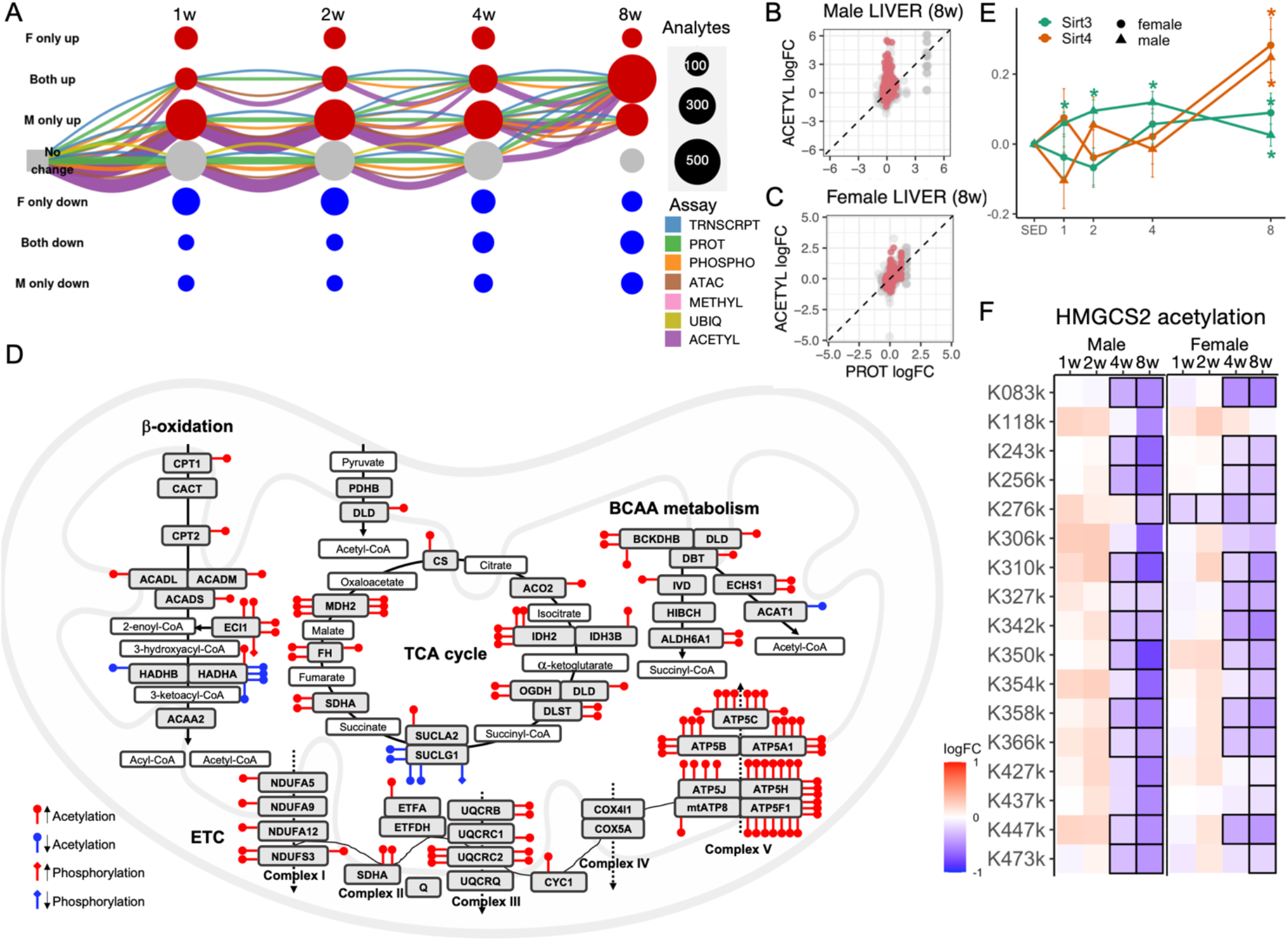
Training-induced mitochondrial adaptation in the liver through protein acetylation. **A)** Graphical representation of the mitochondria-associated training-differential analytes in the liver. Each node represents one of nine possible states (row labels, with F for females and M for males, seven states shown) at each of the four sampled training time points (column labels). Edges are drawn between nodes to represent the path of differential analytes over the training time course, with edge representing the –ome. This graph includes the five largest paths for liver. Both node and edge size are proportional to the number of analytes represented by the node or edge. **B-C)** Correlation between changes (log2 fold change) in protein level and acetylation level in **B)** male and **C)** female liver. Pink color indicates MitoCarta proteins, while other proteins are shown in grey. **D**) Significant acetylation and phosphorylation changes (FDR<0.05) of mitochondrial metabolic proteins in male and female liver after 8 weeks of endurance training (sites changing in only one sex are not illustrated). Each lollipop represents a specific acetylation (rounded top) or phosphorylation (diamond top) site, where red color indicates increases and blue decreases. Multiple lollipops on the same protein indicates several sites significantly changed with training. Proteins with more significant differential sites than could be fitted into the illustration due to space were; Atp5c 9 sites, Atp5a1 9 sites, Atp5h 13 sites, and Idh2 15 sites in total. Proteins displayed with a name different than the official gene name are Atp5b = Atp5f1b, Atp5c = Atp5f1c, Atp5a1 = Atp5f1a, Atp5j = Atp5pf, Atp5h = Atp5pd, Atp5f1 = Atp5pb. **E)** Protein expression changes (log2 fold change) in Sirt3 and Sirt4. Females are represented by circles and males by triangles. * indicates significant change with training (FDR<0.05). **F)** Site-specific acetylation changes in HMGCS2 in males (left panel) and females (right panel). All displayed sites were differentially acetylated overall (taking all timepoints and sexes into account, FDR<0.05), and sites that reach timewise significance (FDR<0.05) are highlighted with black frames.

Notably, in Gene Set Enrichment Analysis (GSEA) that was independent of our graphical results above, males showed transient transcriptional adaptations for multiple pathways that did not result in elevations by week 8 (Fig. S10C). In contrast, in females, the same pathways tended to be upregulated exclusively in week 8. Unlike the transcriptomic data, training-induced changes in mitochondrial proteins were more uniform across males and females (Fig. S10D). A notable sex-consistent exception was observed for mitochondrial dynamics pathways at week 8: these were reduced at the transcription level but elevated at the protein level (Fig. S10D). Recent studies showed upregulation of mitophagy flux in the liver in response to acute exercise^49^ but lower mitophagy/mitochondrial dynamics following training^50^. Therefore, transient increases in mitophagic flux following acute exercise may enhance mitochondrial quality and function after chronic training.

Mitochondrial protein acetylation accounted for over 60% of the significant changes observed in liver following training (Fig. 2A). It is estimated that 35% of mitochondrial proteins are regulated by acetylation and up to 60% of all mitochondrial proteins have acetylation sites^43,51^. The majority of the differentially acetylated mitochondrial proteins remained unchanged for protein abundance (Fig. 6B-C), demonstrating a specific post-translational regulatory mechanism in response to training. We observed significant changes in acetylation of multiple sites across many mitochondrial proteins involved in all major bioenergetic processes (Fig. 6D). In particular, there was an increased acetylation of multiple enzymes involved in lipid transport, lipid catabolism, and oxidative phosphorylation. In addition, increased acetylation was observed for a majority of enzymes in the BCAA degradation pathway, with a concomitant increase in the SLC25A44 protein, a mitochondrial BCAA transporter that increases with training also in human skeletal muscle^52^. Increased acetylation was especially notable in Complex V proteins (Fig. 6D, S10E). Interestingly, increased acetylation of Complex V with exercise training has also been observed in human skeletal muscle^53^, supporting the relevance of our findings for human mitochondrial adaptation. Acetylation of mitochondrial proteins is an important mechanism for regulating fatty acid metabolism^54^, as well as Complex I activity^55^, although the effect of site-specific changes on enzymatic activity remains largely unknown. Acetylation occurs primarily through non-enzymatic mechanisms, i.e. mass action^51,56^, while the deacetylation process occurs through protein deacetylating enzymes. There was a small but significant overall increase in SIRT3 protein across all training timepoints in males, and after 8 weeks also in females (Fig. 6E). These data suggest that greater acetylation of hepatic mitochondrial proteins occurred despite concomitant elevation in SIRT3 activity, similar to the effect observed following training in cardiac tissue. Another mitochondrial sirtuin, SIRT4, also increases in the liver with training.

Next, we delineated training-induced, site-specific acetylation changes of critical hepatic metabolic enzymes. HMGSC2 is the rate-limiting enzyme in the synthesis of ketones. Exercise increases ketogenic flux particularly if performed in postabsorptive conditions^57^; however, the chronic effects of exercise on ketogenic capacity are less known. Liver ketone body production provides an important alternative fuel source during exercise to muscle, heart, and neuronal tissues when glucose levels are challenged, and has been shown to attenuate skeletal muscle proteolysis during high-intensity exercise^58^. We observed deacetylation of lysines 310, 447 and 473 in HMGCS2 (Fig. 6F), which are known to enhance HMGCS2 enzyme activity^59^.

Deacetylation of pyruvate dehydrogenase complex component PDHA1 was also observed (Fig. S10F), and this protein modification enhances enzyme activity^60^, thus promoting oxidative phosphorylation. In addition, acetylation of NDUFS3 (NADH:ubiquinone oxidoreductase core subunit S3 or Complex I 30kDa subunit) and CYP27A1 (bile acid synthesis), increased at multiple sites in the liver after 8 weeks of training. Acetylation of both proteins increased to a greater extent in males (Fig. S10G and H). These findings demonstrate the plasticity of the hepatic mitochondrial proteome and acetylome in response to endurance exercise training and provide novel targets for further mechanistic studies.

### Training upregulates protein networks that are downregulated in type 2 diabetes and cirrhosis in humans

Mitochondrial dysfunction is a hallmark of chronic diseases, including obesity, type 2 diabetes (T2D), nonalcoholic fatty liver disease (NAFLD) and neurodegenerative diseases. We examined the translational relevance of our identified mitochondrial gene sets. First, we performed disease ontology enrichment analysis, identifying disease terms in our training response such that the overlapping disease genes were enriched for MitoCarta genes (see Methods). The heart analytes that were downregulated in response to endurance training (5% FDR) were significantly associated with hypertension, whereas the upregulated analytes were enriched for myopathy terms (Supplementary data S4). The latter was also observed in the skeletal muscle.

Second, we compared our results to nine proteomics disease datasets, which covered skeletal muscle of T2D patients; liver of NAFLD, NASH (nonalcoholic steatohepatitis) and cirrhosis patients; cardiac muscle of hypertrophic cardiomyopathy (HCM) patients; and rodent models of myocardial infarction and heart failure. We focused on proteomics as the majority of the observed changes in these tissues in our study occurred at the protein and protein modification levels. For 8 studies we obtained both differential proteins and the background set of all quantified proteins. When comparing differential proteins in disease to those that were differential in response to exercise, we found significant overlap in T2D for skeletal muscle, obesity and cirrhosis for liver, and HCM and heart failure for cardiac muscle (Fig. 7A). In contrast, there was no significant overlap with protein changes in response to myocardial infarction or NASH.

**Figure 7.**
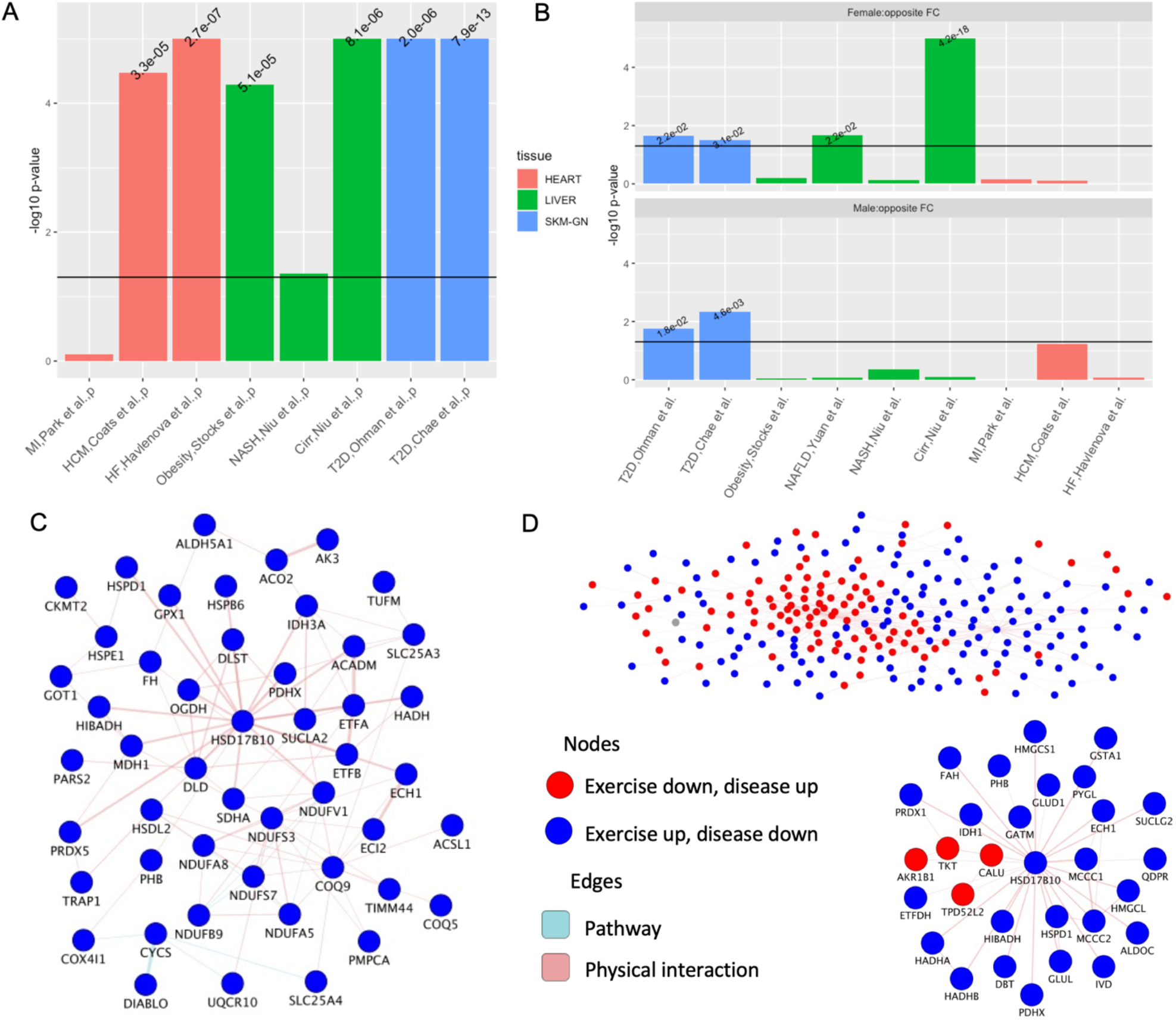
Training results in opposite regulation of mitochondrial proteins compared to type II diabetes and cirrhosis. **A)** Significance of the overlap between the exercise-regulated differential proteins compared to identified proteins in case-control proteomics disease cohorts. The horizontal line represents p=0.05. MI = Myocardial Infarction, HCM = Hypertrophic Cardiomyopathy, NASH = Non-alcoholic Hepatosteatosis, Cirr = Cirrhosis, T2D = Type 2 Diabetes. **B)** Significance of the opposite directionality (Fisher’s exact test) when comparing the fold change sign of the overlapping proteins from (A). NAFLD = Non-alcoholic Fatty Liver Disease, HF = Heart Failure. **C)** Skeletal muscle T2D network. GeneMANIA network of the differential proteins that had sex-consistent response in week 8 of training and were both significant and had opposite direction of effect in two separate T2D cohorts. **D)** Liver cirrhosis network. GeneMANIA network of the 8-week female differential proteins that were both significant and had opposite direction of effect in the liver cirrhosis cohort.

Next we interrogated the directionality of the proteomic signatures in response to training compared to disease. We found a robust, sex-consistent, opposite regulation in response to training in our rats compared to protein signatures of human patients with T2D in two separate datasets (Fig. 7B). Focusing on our sex-consistent differential proteins that were regulated in the opposite direction in both T2D studies, we identified a dense protein-interaction network of exercise-upregulated proteins (Fig 7C). This network includes proteins of the mitochondrial matrix and proteins that are involved in ATP production through the TCA cycle and electron transport chain (q-value <5.4×10^−7^). Moreover, this network contains HSD17B10 (17-beta-hydroxysteroid dehydrogenase 10) as a main hub. This protein is a mitochondrial dehydrogenase involved in fatty acid, amino acid and steroid metabolism. Interestingly, mutations in HSD17B10 cause the mitochondrial disease HSD10, leading to neurodegeneration and cardiomyopathy^61^.

In the liver, there was a striking opposite regulation in cirrhosis patients, uniquely in comparison to the training response in females. The sexual dimorphism in the liver response in this context is relevant considering that men are twice as likely to die from liver cirrhosis compared to women^62^. We overlaid the common differential proteins that had an opposite fold change in exercise compared to disease on known protein-interaction and pathway networks. Exercise-upregulated proteins clustered mostly separately from the exercise-downregulated proteins (Fig 7D). This dichotomy was also observed when performing enrichment analysis: mitochondrial matrix and metabolic pathways were upregulated in response to training (and downregulated in cirrhosis), and RNA processing pathways were instead downregulated in response to training (q-value <3.7×10^−7^). Strikingly, HSD17B10, which was also observed in the T2D network, is the main hub of the exercise-upregulated proteins. In summary, our translational approach reveals multiple candidate mitochondrial proteins and pathways, with HSD17B10 as the main hub gene, for prevention of mitochondrial dysfunction in tissues critical for maintenance of whole-body metabolic health.

## Discussion

While endurance exercise has been known to robustly increase mitochondrial mass and function in skeletal muscle for over 50 years^14^, its impact across other tissues in the body, and the time course of those changes, remains inadequately understood. Here, we provide a landscape of the time-course of mitochondrial multi-omic responses to chronic endurance training across 19 tissues in males and females, reflecting an integrative effect on whole-body metabolism with translational implications for type 2 diabetes and liver cirrhosis. We show that mitochondrial adaptation is largely tissue-, sex- and time course-specific, and we identify major acetylation changes at specific mitochondrial proteins in response to training in the heart and liver, suggesting a novel molecular mechanism for the cardio- and hepato-protective effects of exercise.

Exercise is a powerful modulator of mitochondrial function and health in muscle^16^ and liver^17^, an effect that is conserved across mouse, rat and human. Importantly, while concerted efforts have been made to understand the mitochondrial multi-omic response in one or two tissues^63,64^, this has not been undertaken at the scale or breadth of this work. We show that endurance exercise training activates a concerted mitochondrial response in multiple tissues, including the scarcely investigated colon and adrenal glands. We observe the greatest multi-omic mitochondrial response in skeletal muscle, heart, liver, colon, adrenal gland, BAT, WAT-SC and blood.

Importantly, identifying induction of Complex I phosphorylation as a potential mechanism for training-induced increases in ETC activity in skeletal muscle demonstrates how the breadth and scale of our data allows inspection of where regulation checkpoints may occur. We detected minimal mitochondrial changes in several tissues, including the brain, small intestine and spleen. While brain activity increases during exercise^65^, the brain regions assayed here are already rich in mitochondria with a high rate of glucose consumption at rest^66^ and the increase in metabolic demand during exercise is comparatively low^67^. Exercise-induced vasoconstriction of the renal and splanchnic vasculature^67^ could explain the lack of training-related mitochondrial changes in tissues such as the kidneys, spleen and small intestine.

Despite many common metabolic functions of mitochondria across tissues, the majority of the responses were tissue-specific. While we made initial steps here, further work is required to delineate the specific mechanisms underlying this diversity in response. Potential contributing factors include blood flow, exposure to exerkines, and energy demand and utilization.

Specifically, blood flow to adipose tissue, the contracting muscle and the heart increases during exercise^68^, while flow to the liver is decreased although oxygen uptake increases^69^, with such repeated changes likely driving a mitochondrial response.

We focused on potential mediation through changes in mitochondrial quantity to improve interpretability. Importantly, improvements in quantity (volume) and quality (structure and function) both contribute to overall mitochondrial health and function. We found that the majority of the multi-omic responses were robust to adjustment for markers of mitochondrial volume. Nevertheless, the transcriptomic changes in BAT and the adrenal gland were closely associated with changes in mitochondrial volume. Thermogenesis is the principal function of BAT, a tissue rich in mitochondria^70^. In contrast to females, males lost WAT-SC mass with endurance training^19^. In this context, our observed reduction in the metabolic/thermogenic activity of BAT could serve as an overall energy preservation effect in males. Decreased BAT mass and activity is also observed in chronically trained endurance athletes^71^, corroborating this hypothesis The physiological implications of reduced mitochondrial capacity in the adrenal gland in response to training remains elusive. However, the mitochondria are important stress response modulators known to affect sympathetic adrenal-medullary activation, catecholamine and cortisol/corticosterone levels^72^, and this mitochondrial adaptation is potentially reflective of reduced physiological stress after adapting to repeated bouts of exercise.

Our study has some limitations that should be addressed by future studies. Our use of rat as a model was beneficial for exploring many tissues from the same animals, but required extrapolation to human for the translational analyses. Although we assess several biomarkers of mitochondrial volume, we lack direct functional measurements of mitochondrial respiration in this study. In addition, the acute, transient changes that govern mitochondrial dynamics are not captured using our experimental design, and therefore some of the acute signals that stimulate mitochondrial changes are not detected in this analysis. We are also studying whole organs with multiple cell types that may change in cellular composition following training and responses in specific cell types may be missed. In skeletal muscle, for example, there are increases in the endothelial cell population due to higher vascularization of trained muscle, and changes in fiber type distribution^73^. Finally, a general challenge of mitochondrial adaptation studies is that multiple proteins that localize to mitochondria (48%) can also be found in other cellular compartments^74^. While we addressed these issues indirectly by mechanistic interpretation of our results, deconvolving these two processes systematically poses an interesting computational challenge.

Mitochondrial biogenesis and function are strongly impacted by sex^75^. For example, sexual dimorphism in the skeletal muscle transcriptome at baseline indicate higher lipid oxidation in female skeletal muscle^76^, but sex-specific responses to training are less clear, although the inclusion of more females in exercise studies indicate sex-specific responses^77^. This work provides a foundational resource to understand the importance of sex on temporal mitochondrial adaptations to exercise across 19 different tissues. To this point, we find large sex differences in the temporal dynamics of mitochondrial analytes in response to training in the adrenal gland and BAT. We observed sex-consistent mitochondrial responses in skeletal muscle, as well as sex-specific metabolic responses, including greater induction of lipid oxidation enzymes in females.

Liver mitochondria play a critical role in oxidizing fat (abundant substrate), which provides ATP and substrates to fuel TCA cycle flux and gluconeogenesis^78^ which in turn maintain circulating glucose levels (limited substrate) during fasting and exercise^79^. Thus, hepatic mitochondria serve as a critical energy converter to maintain systemic energy metabolism. Endurance exercise training leads to functional improvements in hepatic mitochondria independent of increases in mitochondrial volume^47,80^, but the molecular mechanisms behind these changes are incompletely understood. Moreover, females show increased hepatic mitochondrial capacity both at baseline and following exercise^50,81^, changes likely due to metabolic demands of gestation and lactation. While it is well-established that chronic exercise training increases protein acetylation in skeletal muscle^82^, much less is known in the heart and liver. We find a dramatic remodeling of the liver mitochondrial acetylome with training, likely due to increased substrate flux through the liver and associated turnover of acetyl CoA in the mitochondria.

Current dogma suggests that nutrient excess causes increased acetylation of mitochondrial proteins that lead to downregulated oxidation metabolism^83^. However, exercise is known to protect hepatic metabolic health and improve oxidative capacity and our results also show dramatic increases in acetylation in response to training. Moreover, exercise training also increases protein acetylation in skeletal muscle^53,82^, which is attributed to improvements in mitochondrial function following training. We observed increased acetylation of mitochondrial proteins in both liver and heart despite a concomitant increase in SIRT3, a mitochondria-specific deacetylase. This finding suggests that SIRT3 is not the cause of the changes in acetylation but may be increasing to control and break a post-translational modification mechanism, such as the shift of the ratio between NAD+ and NADH linked to cellular oxidative stress^84^. Whether acetylation of metabolic proteins plays a functional role in the training-induced improvement in metabolic health is unknown and warrants further study.

Maintaining mitochondrial function is critical for prolonged healthspan, as mitochondrial dysfunction is a hallmark of aging and is associated with cardiovascular, metabolic and neurodegenerative diseases^85^. In contrast, endurance exercise training is a promising intervention for prevention or attenuation of mitochondrial decline. The mitochondrial proteome robustly responded to training in cardiac, skeletal muscle, and liver tissues, which motivated our comparison of the effects in these tissues with proteomic datasets from disease cohorts. We found opposite proteomic skeletal muscle response to training compared to changes observed in human skeletal muscle in T2D patients. There was also opposite regulation of the liver proteome in response to exercise in females compared to changes induced by liver cirrhosis in human patients. Interestingly, the same protein, HSD17B10, was identified as the central hub from protein interaction network analysis of the oppositely regulated proteins in both T2D and liver cirrhosis, though its first degree (exercise) upregulated neighbors were different. HSD17B10 provides a promising target for future therapeutic and mechanistic studies of the health benefits of exercise in skeletal muscle and liver.

Collectively, our work expands upon previous findings by providing an unprecedented multiomic resource of the mitochondrial adaptation to endurance exercise training. We have concurrently mapped mitochondrial changes to endurance exercise in 19 different tissues. Importantly, this MoTrPAC resource provides much-needed insight into sex-specific mitochondrial adaptations and how these changes occur over time. Altogether, considering the critical role that mitochondria play in maintaining tissue-specific and whole body metabolic health, this work provides an unparalleled resource to stimulate hypothesis-driven, mechanistic studies, as well as work aimed at identifying targets that can be leveraged therapeutically to combat mitochondrial dysfunction and metabolic diseases.

## Methods

### Experimental design

#### Exercise training protocol

Inbred male and female Fischer 344 rats were obtained from the National Institute of Aging (NIA) rodent colony. Rats were housed in pairs at a reverse dark-light cycle, kept at a temperature of 20-25°C and fed normal chow (Lab Diet 5L79). After familiarization (>10 days for reverse light cycle, 12 days for treadmill), rats were randomized to training or control. The rats were partitioned into three groups; 8-week rats that were randomized to training or control, 4-week rats that were all assigned to training, and 1- and 2-week rats that were randomly assigned to 1- or 2-weeks of training. A total of 50 rats (5 males and 5 females per time point) were used for molecular analyses, with the exception of proteomics that was performed on 60 animals (6 males and 6 females per time point). All training groups started training at 6 months of age and trained on a Panlab 5-lane rat treadmill (Harvard Instruments, Model LE8710RTS). Rats were exercised 5 days per week using a progressive protocol aimed to maintain an intensity corresponding to approximately 70% of VO_2_max (increasing grade and speed, see^19^ for details), with a maximal duration of 50min for the last two weeks of training. The starting treadmill speed was based on VO_2_max measurements obtained following familiarization. All animal procedures took place during the dark cycle and were approved by the Institutional Animal Care and Use Committee at the University of Iowa, where the training intervention took place.

#### Phenotyping

Body composition was measured using nuclear magnetic resonance (Minispec LF90II Body Composition Rat and Mice Analyzer) for all rats prior to training, and for the 4-week and 8-week animals five days prior to tissue harvest. Maximal oxygen consumption (VO_2_max) was similarly measured in all rats prior to training, and during the last week of training for the rats in the 4-week and 8-week training groups and the sedentary group. The testing protocol consisted of a 15-minute warm up at a treadmill speed of 9 m/min and 0° incline. The incline was subsequently increased to 10° and the speed was increased by 1.8 m/min every 2 minutes^86^ until exhaustion, defined as when the rat sat on the shock area 3 consecutive times without responding to increased shock. Blood for lactate assessment was taken from the tail immediately after the test. The criteria for reaching VO_2_max was a leveling off of VO_2_ despite increased workload, a respiratory exchange ratio >1.05, and a blood lactate concentration ≥6 mM.

#### Tissue collection

All tissues were collected 48 hours after the last exercise bout. Food was removed three hours prior to the start of dissections, for which rats were sedated with inhaled isoflurane (1-2%) and kept under anesthesia until death. Blood was obtained through cardiac puncture, then the gastrocnemius muscle, subcutaneous white fat, right lobe of the liver, heart, and lungs were removed in that particular order. Removal of the heart resulted in death. A guillotine was subsequently used for decapitation, after which the brain was removed, and the hypothalamus, right and left hippocampus, right and left cerebral cortex were dissected out. After decapitation, the right kidney, right and left adrenal glands, spleen, brown adipose tissue, small intestine, colon, right testes or ovaries and right vastus lateralis were removed in that order. All tissues were flash-frozen in liquid nitrogen and stored at -80°C.

#### Data generation and processing

Aliquots of assay- and tissue-specific reference standards were included in all molecular assays to evaluate technical differences across batches. All assays, omic quantification pipelines, and quality assurance processes are described in^19^ and an overview is provided in the supplementary text.

#### Reannotation of cardiolipin data

In comparison to the initial dataset^19^, cardiolipin data was re-annotated after improvements to the spectral libraries were introduced. We revisited the annotations to capture more compounds in the class across tissues compared to the initial dataset.

#### mtDNA quantification

The protein/DNA precipitate resulting from the organic extraction for metabolomics was dried under vacuum for one hour in a GeneVac EZ-2 evaporator. Dried samples were stored at -20°C until DNA extraction. DNA was extracted from dried pellets using a Qiagen DNeasy Blood and Tissue kit (Qiagen, Germany, #69506). The DNA concentration in the eluate was determined and 50 ng of sample DNA was used for multiplex qPCR Mitochondrial levels were quantified following the protocol of Nicklas et al^87^ using a 5’VIC reporter and a 3’TAMRA quencher dye and D-loop expression with a 5 ‘6-FAM reporter and 3’ TAMRA-labeled quencher. Amplification was carried in a 25ul reaction consisting of 1x TaqMan Universal Master Mix II (ThermoFisher, MA), using 200 nM each β-actin forward (GGGATGTTTGCTCCAACCAA) and reverse primers (GCGCTTTTGACTCAAGGATTTAA) to estimate nuclear DNA and 50 nM each mitochondrial D-loop forward (GGTTCTTACTTCAGGGCCATCA) and reverse (GATTAGACCCGTTACCATCGAGAT) primer and 100 nM each B-actin probe (VIC-CGGTCGCCTTCACCGTTCCAGTT-TAMRA) and D-loop probe (6FAM-TTGGTTCATCGTCCATACGTTCCCCTTA-TAMRA). Forty cycles of amplification were performed on duplicate samples and relative mitochondrial levels calculated as C_T_(mito) - C_T_(nuclear) using the 2^−ΔΔCT^ method^88^.

#### Statistical analyses

##### Differential analysis and mitochondrial analyte selection

Differential analysis was conducted in each tissue, -ome and sex separately. The full details on the processing has been described elsewhere^19^. Briefly, DESeq2^89^ was used for RNA-Seq, edgeR^90^ for RRBS, and limma^91^ for proteomics, metabolomics, and ATAC-seq data. As input, normalized data were used for proteomics, and ATAC-seq, and filtered raw counts were used for RNA-seq, and RRBS. For targeted metabolomics, the KNN-imputed (if it included > 12 analytes) log_2_-transformed data were used; otherwise, log_2_-transformed data were used. For untargeted metabolomics, log_2_ KNN-imputed data were used. F-tests (limma, *edgeR::glmQLFTest*) or likelihood ratio tests (*DESeq2::nbinomLRT, lrtest*) were used to identify analytes that changed over the 8-week training time course. Male- and female-specific p-values were combined using Fisher’s sum of logs meta-analysis into a single p-value (*training p-value*) and p-value adjustment was performed using Independent Hypothesis Weighting (IHW)^92^, with IHW FDR ≤ 5% for each ome across tissues. Time-and sex-specific effects were calculated by comparing each training time point with its sex-matched sedentary control animals (*timewise summary statistics*) using the following functions: *limma::contrasts*.*fit* with *limma::eBayes, DESeq2::DESeq* and *edgeR::glmQLFTest*. Assay-specific covariates were included based on technical metrics (RNA integrity number, median 5’-3’ bias, percent of reads mapping to globin, and percent of PCR duplicates as quantified with Unique Molecular Identifiers (UMIs) for RNA-Seq; fraction of reads in peaks and library preparation batch for ATAC-seq). For metabolomics data, meta-regression of the 1116 metabolites was performed using R’s *metafor* package^93^.

MitoCarta 3.0^3^ was used to select mitochondria-associated genes (RNAseq, RRBS and ATAC-seq) and proteins (global proteomics, phosphoproteomics, acetylome and ubiquitylome). The MitoCarta list included 1136 human and 1140 mouse genes identified through mass-spectrometry on isolated mitochondria from 14 different tissues, in combination with GFP-tagging for localization, integration with other datasets and literature curation. This resource also assigns genes to a mitochondria-specific ontology, the MitoCarta 3.0 mitochondrial pathways (149 in total)^3^. The rat ortholog mapping of MitoCarta that was used to select mitochondrial IDs is provided in Supplementary data S7. Metabolite selection was based on Heden et al.^24^, see Table S5 for the full list of metabolites used.

##### Graphical clustering analysis

Graphical clustering analysis of the timewise summary statistics was described in^19^. In this work the graphical representation results were filtered down to represent the MitoCarta analytes (e.g., for Fig. 3C, Supplementary data S3). We now briefly describe this graphical method. Z-scores of the IHW-selected analytes were modeled using a mixture distribution to separate null from non-null cases and identify clusters while accounting for correlation over time and between the sexes. Let *Z* ∈ *R*^*n x t x 2*^represent the input, where z_i,j,k_ represents the z-score of analyte *i* ∈ {1,…,n} at the training time point *j* ∈ {1,…,t} of sex *k* ∈ {m,f}, with m=males and f=females. Under the assumption that z_i,j,k_ follows a mixture distribution of null and non-null z-scores, each z_i,j,k_ has a latent configuration h_i,j,k_ ∈ {-1,0,1}, where -1 denotes downregulation, 0 denotes null (no change), and 1 denotes upregulation. A *full configuration matrix* (e.g., specifying if a z-score is null, up, or down for each time point in each sex) is denoted **h** ∈ {−1,0,1}^t *x 2*^, and **z**_i_ ∈ R^t *x 2*^ is the matrix of all z-scores of analyte *i*. The expectation-maximization (EM) process of the repfdr algorithm^26,94^ was used to estimate for each possible **h** both its prior probability π(**h**) and its posterior Pr(**h**|**z**_i_), for every analyte *i*. The *locfdr* R package ^95^, is used in this process to infer the marginal mixture distribution of each time point *j* and sex *k*. That is, all z-scores (i.e., not limited to mitochondrial analytes) z_*jk_ are used to estimate the densities: *f*_*j,k*_*(z*|*H*_i,*j,k*_ = −1) = *f*_*+,*j,k*_*(z*), *f*_*j,k*_*(z*|*H*_i,*j,k*_ = 0) = *N(*0,1), and *f*_*j,k*_*(z*|*H*_i,*j,k*_k= −1) = *f*_+,*j,k*_*(z*). We excluded configurations **h** with π(**h**) < 0.001 and normalized Pr(**h**|**z**_i_) to sum to 1. The new posteriors can be interpreted as a soft clustering solution, where the greater the value is, the more likely it is for analyte *i* to participate in cluster **h**.

We use these posteriors to assign analytes to “states”, where a state is a tuple (s_m,j_, s_f,j_), where s_m,j_ is the differential abundance state null, up, or down (0,1, and -1 in the notation above, respectively) in males at time point *j* (s_f,j_ corresponds to females at time point *j*), resulting in nine possible states in each time point. For example, assume we inspect analyte *i* in time point *j*, asking if the abundance is upregulated in males while null in females, then we sum over all posteriors Pr(h|z_i_) such that h_m,j_=1 and h_f,j_=0. If the result is greater than 0.5, then we assign analyte *i* to the node set S(s_m,j_, s_f,j_). We use S(s_m,j_, s_f,j_) to denote all analytes that belong to a state (s_m,j_, s_f,j_) and for every pair of states from adjacent time points *j* and *j+1* we define their edge set E(s_m,j_, s_f,j_, s_m,j+1_, s_f,j+1_) as the intersection of S(s_m,j_, s_f,j_) and S(s_m,j+1_, s_f,j+1_). Note that these can be defined using a similar marginalization as was done to define the node sets, but in practice we found that these two approaches resulted in almost identical results. The sets S and E together define a tree structure that represent the differential patterns over sex and time.

##### Enrichment analyses and pathway annotation of selected sets

Analytes were mapped to Ensembl gene IDs. For each identified analyte set (e.g., a node or an edge set from the graphical clustering above), we performed pathway enrichment analysis using the full MitoCarta 3.0 gene list as background. Enrichment analysis was also performed using the KEGG and REACTOME rat pathways (organism “rnorvegicus”) using the *gprofiler2::gost* function in R^96^. Nominal p-values were calculated using a one-tailed hypergeometric test, and were then adjusted across all results using IHW with tissue as a covariate. Pathways with a q-value < 0.1 were considered significant. Fig. 2C displays significantly enriched pathways (TRNSCRPT, PROT, ACETYL) for the 8-week node for all tissues with at least one enriched pathway (<5% BH FDR), the top pathway from each MitoCarta subcategory with the greatest number of enrichments in shown. All enrichment results are available in Supplementary data S6.

To identify which mitochondrial pathways were most differentially regulated within each tissue across timepoints and within each sex, we used our identified repfdr sets (see previous section) together with the MitoCarta 3.0 human database. Here, each significant analyte was annotated to one of 8 major MitoCarta mitochondrial pathway groups including metabolism, mitochondrial central dogma, mitochondrial dynamics and surveillance, oxidative phosphorylation, protein import, sorting and homeostasis, signaling, small molecule transport and “other”. Analytes (proteins or genes) within each pathway were then separated by time point and differential regulation by sex according to state score. For example, for a given time point we can assign analytes to a pattern that represents upregulation in both males and females (denoted as F1_M1), or upregulation in females and downregulation in males (denoted as F1_M-1). This representation was used to display the number of time-dependent and sex-specific differentially regulated features in each pathway.

##### PCA

Principal Component analysis with scaling was performed on the sedentary 8-week control animals from the RNA-seq and proteomics data separately. The first two components contributed to the majority of the variance in both the datasets with the first component revealing clear tissue-specific differences.

##### Biological network analyses

Pathway and protein interaction networks of a selected set of genes were created using GeneMANIA^97^ and visualized using the Cytoscape software^98^.

##### Disease ontology enrichment analysis

We first filtered the disease ontology database before applying the enrichment analyses. Our rationale here was that many disease terms may be enriched with general biological processes that are relevant for many tissues both in health and disease states (e.g., cell proliferation in cancer disease terms), and are thus not likely to reflect a true association between our exercise-specific results and diseases. We therefore generated tissue-specific disease ontology terms by utilizing gene expression data from GTEx v8^99^. For each disease ontology term and a tissue (covered by GTEx) we computed the p-value for the overlap between the term’s gene set and the tissue’s gene set. If the p-value was greater than 0.01 then we omitted the term from the tissue’s analyses. Disease ontology enrichment analysis was then performed using the DOSE R package^98^ for each of our tissue- and ome-specific gene sets that had at least 10 genes. For our mitochondrial-focused analysis we then report results that are: (1) significant at 5% FDR, (2) had at least three genes in the intersection between our set and the disease term gene set, and (3) the overlapping gene set from (2) was significantly enriched for MitoCarta genes (p<0.05, hypergeometric test). The results are available in Supplementary data S4.

##### Gene and post-translational modification set enrichment analyses

Gene set enrichment analysis (GSEA) and post-translational modification set enrichment analysis (PTM-SEA) was performed using ssGSEA2.0^100^. The input for GSEA was the t-scores from the timewise comparisons for all analytes (not just mitochondrial). Here, analytes were integrated into gene ids by taking the most significant t-score (i.e., the one with the maximum absolute value). Phosphosite-level t-scores and the human PTMSigDB^100^ were used as input for PTM-SEA. We used the MitoPathways database from MitoCarta 3.0^3^ to identify enriched mitochondrial pathways. Human gene symbols were mapped to rat orthologs before running the analysis. We used the NCBI Reference Protein Sequence database (RefSeq) to annotate protein IDs, and mapped PTM sites from rats to humans using BLASTp to align rat sequences to the human UniProt fasta sequence database, and used alignments with >60% sequence identity for mapping. For both GSEA and PTM-SEA, we ran the *ssGSEA2* function with parameters that avoid normalization, required at least 5 overlapping features with the gene set, and used the area under the curve as the enrichment metric (sample.norm.type = “none”, weight=0.75, correl.type = “rank”, statistic = “area.under.RES”, output.score.type = “NES”, min.overlap=5).

##### DREM

We used DREM^35^ for network inference of transcription factors driving the transcriptional changes in specific sex and tissue combinations across the 8-week training time course. Selecting for MitoCarta genes, we used the z-score per gene in each time point as input. For transcription factor-target data we used the network inferred by NicheNet^101^. To use the NicheNet network, we mapped the rat gene symbols to human symbols using data from RGD (v39). We used a low penalty for adding nodes (40) and a convergence likelihood of 0.01%. We also ran the models using all the transcriptome results to confirm that the predicted transcription factors were identified using the entire transcriptome as well, with the appropriate background.

##### Comparison to disease datasets

Our differential proteomic results from the endurance training intervention were compared to case-control proteomic results from disease cohorts. The skeletal muscle results were compared to two human skeletal muscle T2D cohorts^102,103^, the liver results were compared to human liver datasets for NASH, cirrhosis^104^ and NAFLD^105^, as well as a mouse dataset on obesity^106^. The cardiac results were compared to a human cardiac proteomic study on hypertrophic cardiomyopathy^107^, and the effects of myocardial infarction^108^ and heart failure^109^ in rat cardiac muscle. For each comparison of rat vs. human results we first subsetted the data to the protein ids shared by both platforms. That is, we set the background for the comparison to the set of proteins that were quantified (i.e., not necessarily significantly differential) in our platforms and that of the compared human study (when available). Then, we computed the significance of the overlap between the human study reported significant protein ids and our IHW-selected protein ids via Fisher’s exact test. Among the proteins that were in this overlap, we again tested for directionality of the effects using Fisher’s exact test, but with the alternative being of overlap lower than expected. For example, if 10 proteins were identified as significant in both the human study and our rat study, then we annotated each one by their sign of differential abundance as up/down (week 8 results from the rat, performed for each sex separately). Then, we applied Fisher’s exact test for the null hypothesis that the sign concordance between the two resources is random, and the alternative that the discordance is greater than expected by chance.

## Abbreviations

Abbreviation: Definition
BAT: Brown Adipose Tissue
BCAA: Branched Chain Amino Acid
DREM: Dynamic Regulatory Events Miner
ETC: Electron Transport Chain
GSEA: Gene Set Enrichment Analysis
HCM: Hypertrophic Cardiomyopathy
IHW: Independent Hypothesis Weighting
mtDNA: Mitochondrial DNA
NAFLD: Nonalcoholic Fatty Liver Disease
NASH: Nonalcoholic Steatohepatitis
OXPHOS: Oxidative Phosphorylation
PCA: Principal Component Analysis
RRBS: Reduced Representation Bisulfite Sequencing
SED: Sedentary control animals
SKM-GN: *Gastrocnemius* Skeletal Muscle
SKM-VL: *Vastus Lateralis* Skeletal Muscle
T2D: Type 2 Diabetes
TCA: Tricarboxylic Acid Cycle
WAT-SC: White Adipose Tissue - Subcutaneous

## Funding

The MoTrPAC Study is supported by NIH grants U24OD026629 (Bioinformatics Center), U24DK112349, U24DK112342, U24DK112340, U24DK112341, U24DK112326, U24DK112331, U24DK112348 (Chemical Analysis Sites), U01AR071133, U01AR071130, U01AR071124–01, U01AR071128, U01AR071150, U01AR071160, U01AR071158 (Clinical Centers), U24AR071113 (Consortium Coordinating Center), U01AG055133, U01AG055137 and U01AG055135 (PASS/Animal Sites).

Parts of this work were performed in the Environmental Molecular Science Laboratory, a U.S. Department of Energy national scientific user facility at Pacific Northwest National Laboratory in Richland, Washington.

## Data and code availability

All -omics data from this study, including training differential results for all tissues and -omes, are available in the R package https://motrpac.github.io/MotrpacRatTraining6moData/.

Timewise differential results adjusted for biomarkers of mitochondrial volume are available as Extended Data S1, see https://zenodo.org/record/7459795#.Y6H4OuzMK-Y (tables are available as RData files). Analysis code for reproducing the results from this study are available here: https://github.com/MoTrPAC/motrpac-rat-training-mitochondria. The cardiolipin and mtDNA data are also available in this repository.

## Supplemental and Extended Data Tables

**Extended Data S1**. Differential analysis adjusted for markers of mitochondrial volume.

**Supplementary data S1**. Training effect on biomarkers of mitochondrial volume.

**Supplementary data S2**. Clustering results for pre- and post-adjustment.

**Supplementary data S3**. Repfdr results selected for mitochondrial analytes.

**Supplementary data S4**. Disease ontology results.

**Supplementary data S5**. List of metabolites used.

**Supplementary data S6**. MitoCarta 8-week enrichment results.

**Supplementary data S7**. MitoCarta rat ortholog list.

