## Supplementary figures for "The mitochondrial multi-omic response to exercise training across tissues"

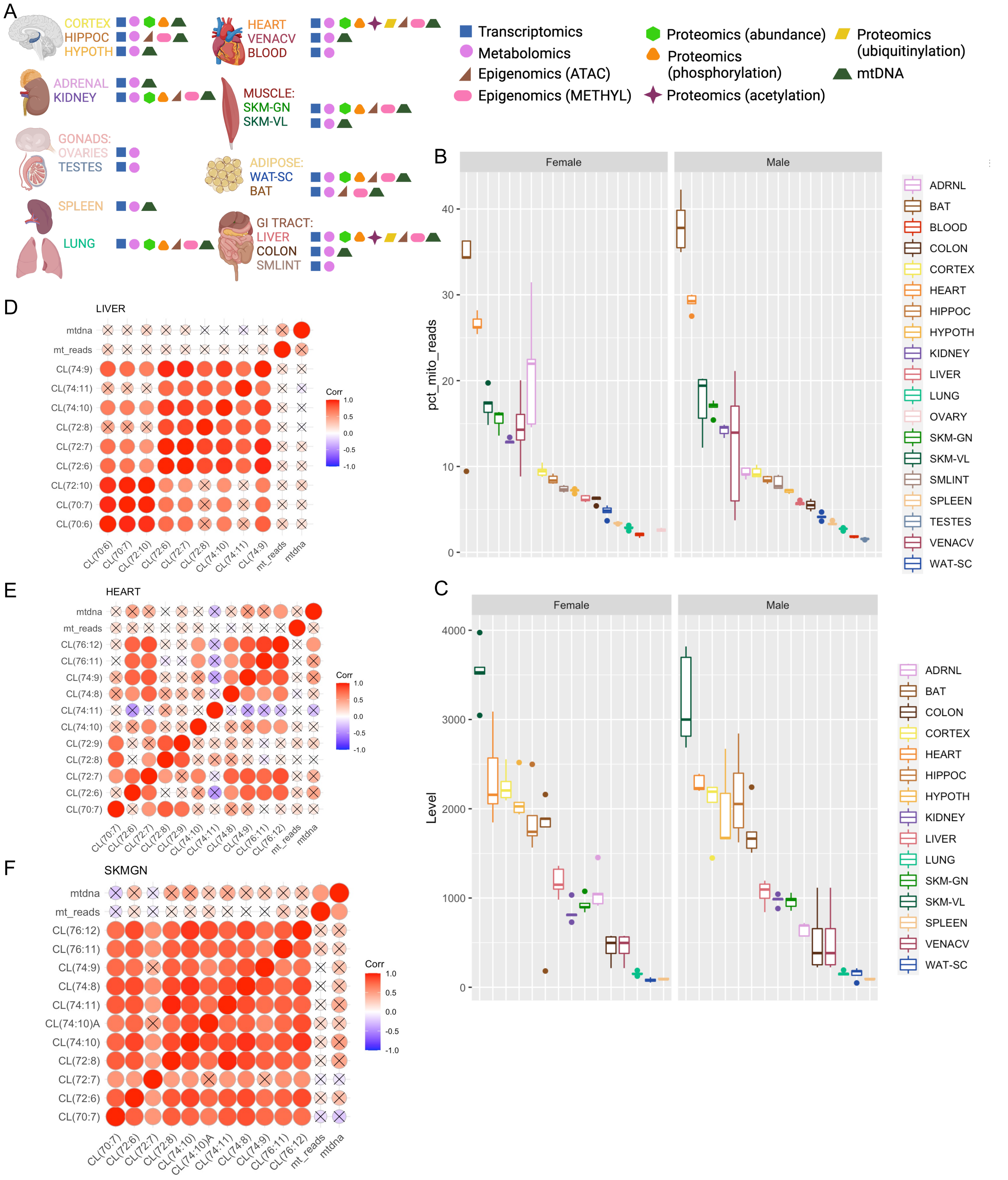
A
CORTEX
HIPPOC
HYPOTH
HEART
VENACV
BLOOD
ADRENAL
KIDNEY
MUSCLE:
SKM-GN
SKM-VL
GONADS:
OVARIES
TESTES
ADIPOSE:
WAT-SC
BAT
SPLEEN
GI TRACT:
LIVER
COLON
SMLINT
LUNG
Transcriptomics
Metabolomics
Epigenomics (ATAC)
Epigenomics (METHYL)
Proteomics (abundance)
Proteomics (phosphorylation)
Proteomics (acetylation)
Proteomics (ubiquitinylation)
mtDNA
B
Female
Male
pct_mito_reads
0
10
20
30
40
ADRNL
BAT
BLOOD
COLON
CORTEX
HEART
HIPPOC
HYPOTH
KIDNEY
LIVER
LUNG
OVARY
SKM-GN
SKM-VL
SMLINT
SPLEEN
TESTES
VENACV
WAT-SC
C
Female
Male
Level
0
1000
2000
3000
4000
ADRNL
BAT
COLON
CORTEX
HEART
HIPPOC
HYPOTH
KIDNEY
LIVER
LUNG
SKM-GN
SKM-VL
SPLEEN
VENACV
WAT-SC
D
LIVER
mtdna
mt_reads
CL(74:9)
CL(74:11)
CL(74:10)
CL(72:8)
CL(72:7)
CL(72:6)
CL(72:10)
CL(70:7)
CL(70:6)
Corr
1.0
0.5
0.0
-0.5
-1.0
E
HEART
mtdna
mt_reads
CL(76:12)
CL(76:11)
CL(74:9)
CL(74:8)
CL(74:11)
CL(74:10)
CL(72:9)
CL(72:8)
CL(72:7)
CL(72:6)
CL(70:7)
Corr
F
SKMGN
mtdna
mt_reads
CL(76:12)
CL(76:11)
CL(74:9)
CL(74:8)
CL(74:11)
CL(74:10)A
CL(74:10)
CL(72:8)
CL(72:7)
CL(72:6)
CL(70:7)
Corr

**Supplementary Figure 1. Molecular assays and biomarker analysis. A)** Overview of assays run for each tissue. HIPPOC = Hippocampus, HYPOTH = Hypothalamus, SMLINT = Small Intestine, SKM-GN = Skeletal Muscle Gastrocnemius, SKM-VL = Skeletal Muscle Vastus Lateralis, WAT-SC = White Adipose Tissue, Subcutaneous, BAT = Brown Adipose Tissue, VENACV = Vena Cava. Created using BioRender.com. **B)** Distribution of percent mitochondrial reads from RNA-seq analysis of all tissues. Each boxplot shows the distribution of the values in the control (untrained) animals. **C)** Distribution of mtDNA levels across the analyzed tissues. Each boxplot shows the distribution of the values in the control (untrained) animals. In both **C** and **D** the whiskers extend from the hinge to the largest and lowest values, but no further than 1.5 * (the interquartile range). **D-F)** Correlation between different biomarkers of mitochondrial volume, including different cardiolipin species in **D)** Liver, **E)** Heart and **F)** Skeletal muscle (*gastrocnemius*). For these plots, Pearson correlations were computed using all animals.

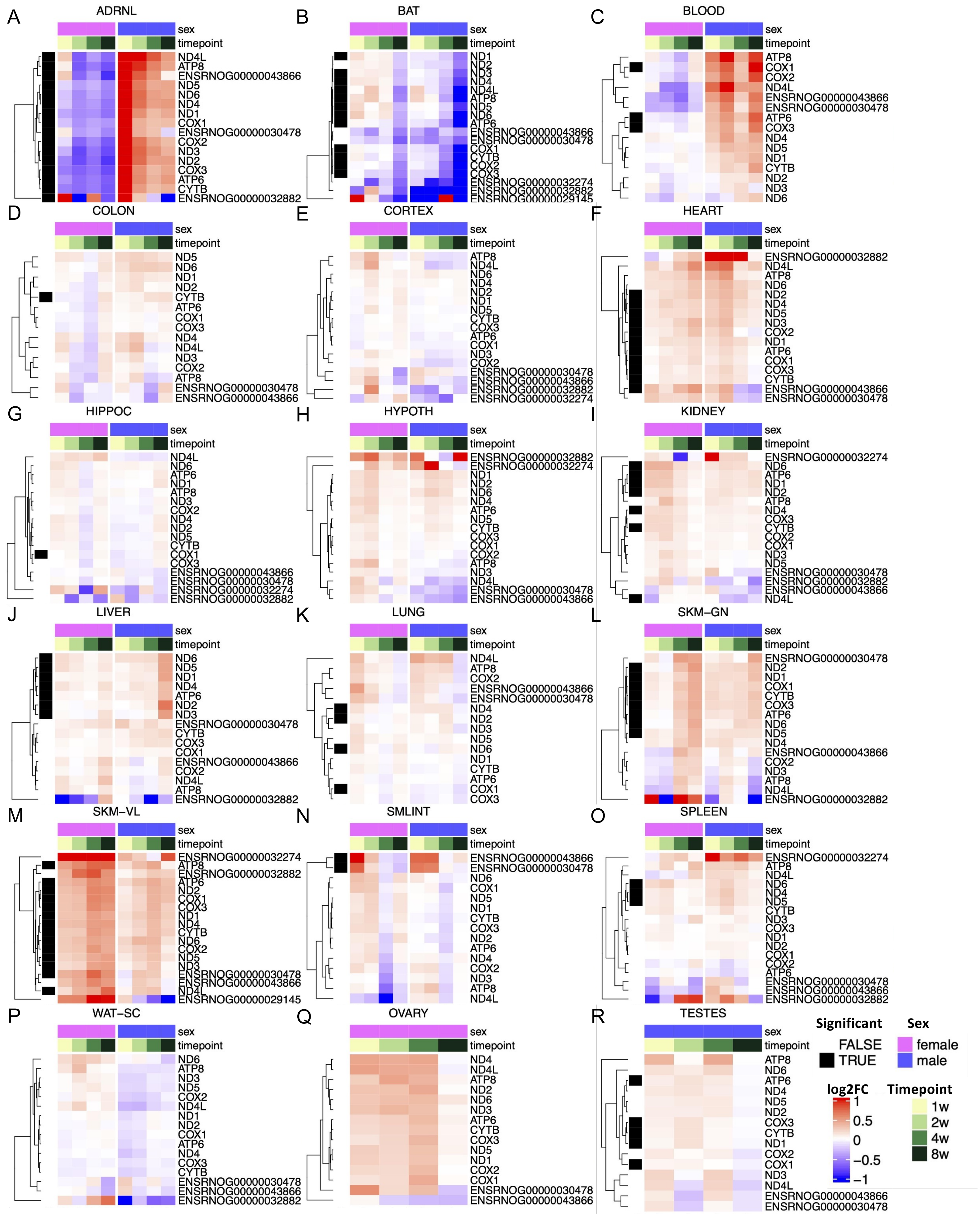
A
ADRNL
sex
timepoint
ND4L
ATP8
ENSRNOG00000043866
ND5
ND6
ND4
ND1
COX1
ENSRNOG00000030478
COX2
ND3
ND2
COX3
ATP6
CYTB
ENSRNOG00000032882
B
BAT
sex
timepoint
ND1
ND2
ND3
ND4
ND4L
ATP8
ND5
ND6
ATP6
ENSRNOG00000043866
ENSRNOG00000030478
COX1
CYTB
COX2
COX3
ENSRNOG00000032274
ENSRNOG00000032882
ENSRNOG00000029145
C
BLOOD
sex
timepoint
ATP8
COX1
COX2
ND4L
ENSRNOG00000043866
ENSRNOG00000030478
ATP6
COX3
ND4
ND5
ND1
CYTB
ND2
ND3
ND6
D
COLON
sex
timepoint
ND5
ND6
ND1
ND2
CYTB
ATP6
COX1
COX3
ND4
ND4L
ND3
COX2
ATP8
ENSRNOG00000030478
ENSRNOG00000043866
E
CORTEX
sex
timepoint
ATP8
ND4L
ND6
ND4
ND2
ND1
ND5
CYTB
COX3
ATP6
COX1
ND3
COX2
ENSRNOG00000030478
ENSRNOG00000043866
ENSRNOG00000032882
ENSRNOG00000032274
F
HEART
sex
timepoint
ENSRNOG00000032882
ND4L
ATP8
ND6
ND2
ND4
ND5
ND3
COX2
ND1
ATP6
COX1
COX3
CYTB
ENSRNOG00000043866
ENSRNOG00000030478
G
HIPPOC
sex
timepoint
ND4L
ND6
ATP6
ND1
ATP8
ND3
COX2
ND4
ND2
ND5
CYTB
COX1
COX3
ENSRNOG00000043866
ENSRNOG00000030478
ENSRNOG00000032274
ENSRNOG00000032882
H
HYPOTH
sex
timepoint
ENSRNOG00000032882
ENSRNOG00000032274
ND1
ND2
ND6
ND4
ATP6
ND5
CYTB
COX3
COX1
COX2
ATP8
ND3
ND4L
ENSRNOG00000030478
ENSRNOG00000043866
I
KIDNEY
sex
timepoint
ENSRNOG00000032274
ND6
ATP6
ND1
ND2
ATP8
ND4
COX3
CYTB
COX2
COX1
ND3
ND5
ENSRNOG00000030478
ENSRNOG00000032882
ENSRNOG00000043866
ND4L
J
LIVER
sex
timepoint
ND6
ND5
ND1
ND4
ATP6
ND2
ND3
ENSRNOG00000030478
CYTB
COX3
COX1
ENSRNOG00000043866
COX2
ND4L
ATP8
ENSRNOG00000032882
K
LUNG
sex
timepoint
ND4L
ATP8
COX2
ENSRNOG00000043866
ENSRNOG00000030478
ND4
ND2
ND3
ND5
ND6
ND1
CYTB
ATP6
COX1
COX3
L
SKM−GN
sex
timepoint
ENSRNOG00000030478
ND2
ND1
COX1
CYTB
COX3
ATP6
ND6
ND5
ND4
ENSRNOG00000043866
COX2
ND3
ATP8
ND4L
ENSRNOG00000032882
M
SKM−VL
sex
timepoint
ENSRNOG00000032274
ATP8
ENSRNOG00000032882
ATP6
ND2
COX1
COX3
ND1
ND4
CYTB
ND6
COX2
ND5
ND3
ENSRNOG00000030478
ENSRNOG00000043866
ND4L
ENSRNOG00000029145
N
SMLINT
sex
timepoint
ENSRNOG00000043866
ENSRNOG00000030478
ND6
COX1
ND5
ND1
CYTB
COX3
ND2
ATP6
ND4
COX2
ND3
ATP8
ND4L
O
SPLEEN
sex
timepoint
ENSRNOG00000032274
ATP8
ND4L
ND6
ND4
ND5
CYTB
ND3
COX3
ND1
ND2
COX1
COX2
ATP6
ENSRNOG00000030478
ENSRNOG00000043866
ENSRNOG00000032882
P
WAT−SC
sex
timepoint
ND6
ATP8
ND3
ND5
COX2
ND4L
ND1
ND2
COX1
ATP6
ND4
COX3
CYTB
ENSRNOG00000030478
ENSRNOG00000043866
ENSRNOG00000032882
Q
OVARY
sex
timepoint
ND4
ND4L
ATP8
ND2
ND6
ND3
ATP6
CYTB
COX3
ND5
ND1
COX2
COX1
ENSRNOG00000030478
ENSRNOG00000043866
R
TESTES
sex
timepoint
ATP8
ND6
ATP6
ND4
ND5
ND2
COX3
CYTB
ND1
COX2
COX1
ND3
ND4L
ENSRNOG00000043866
ENSRNOG00000030478
Significant
FALSE
TRUE
Sex
female
male
log2FC
1
0.5
0
−0.5
−1
Timepoint
1w
2w
4w
8w

**Supplementary Figure 2. The mitochondria-encoded transcriptional changes per timepoint, sex and tissue.** Changes in transcriptional abundance (log2 fold change) for mitochondria-encoded transcripts across the 8 weeks of training (weeks 1, 2, 4 and 8) vs. controls, for each sex (females labeled by a pink bar and males by a blue bar). Significant genes are indicated by a black bar. Data are shown for 18 tissues (vena cava excluded due to fewer samples, see methods); **A)** Adrenal glands, **B)** Brown adipose tissue, **C)** Blood, **D)** Colon, **E)** Cortex, **F)** Heart, **G)** Hippocampus, **H)** Hypothalamus, **I)** Kidney, **J)** Liver, **K)** Lung, **L)** Skeletal muscle (Gastrocnemius), **M)** Skeletal muscle (Vastus Lateralis), **N)** Small intestine, **O)** Spleen, **P)** White adipose tissue, **Q)** Ovary, **R)** Testes.

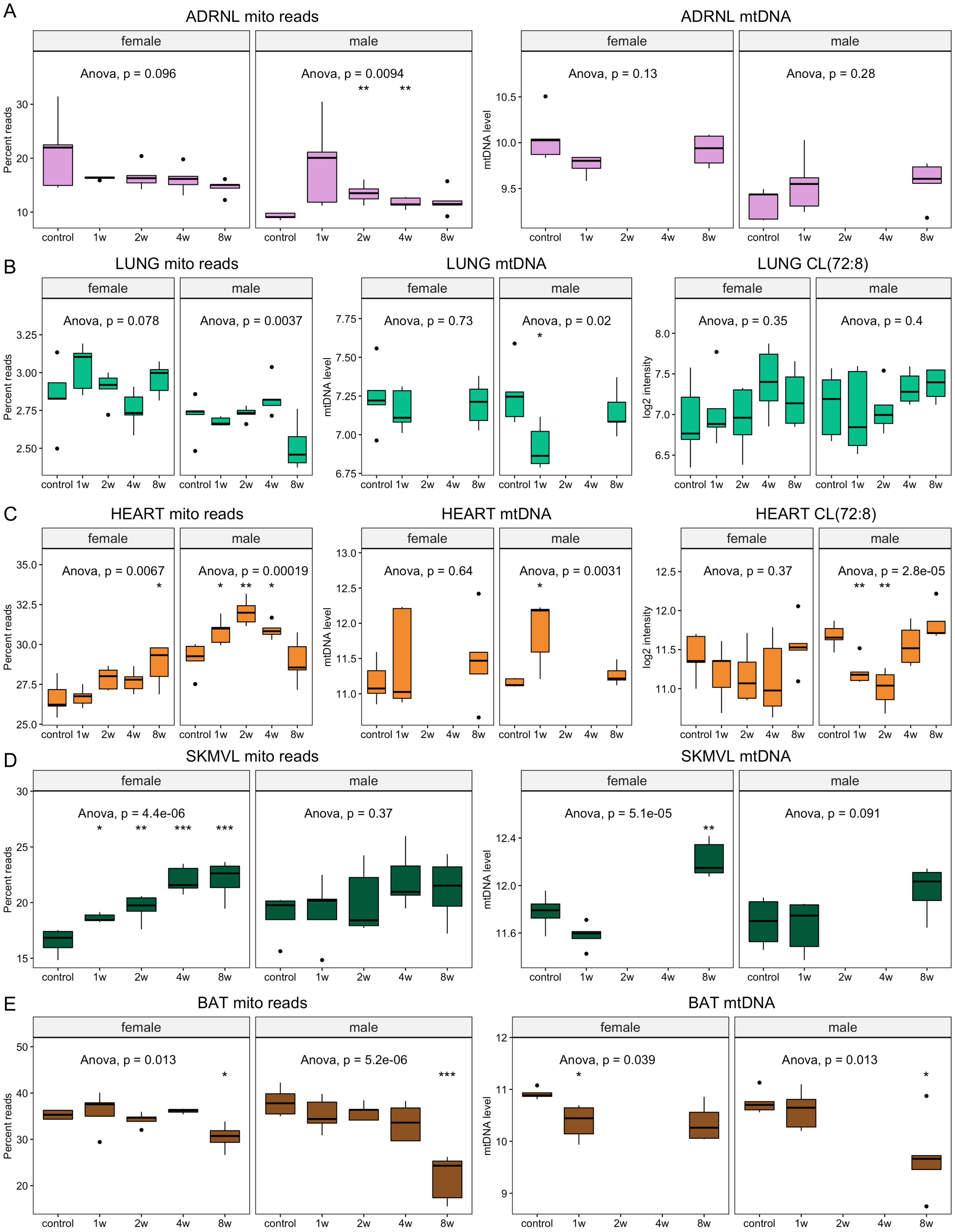
A
ADRNL mito reads
female
male
Anova, p = 0.096
Anova, p = 0.0094
** **
Percent reads
30
20
10
control 1w 2w 4w 8w
ADRNL mtDNA
Anova, p = 0.13
Anova, p = 0.28
mtDNA level
10.5
10.0
9.5
B
LUNG mito reads
Anova, p = 0.078
Anova, p = 0.0037
3.25
3.00
2.75
2.50
LUNG mtDNA
Anova, p = 0.73
Anova, p = 0.02
*
7.75
7.50
7.25
7.00
6.75
LUNG CL(72:8)
Anova, p = 0.35
Anova, p = 0.4
log2 intensity
8.0
7.5
7.0
6.5
C
HEART mito reads
Anova, p = 0.0067
*
Anova, p = 0.00019
* ** *
35.0
32.5
30.0
27.5
25.0
HEART mtDNA
Anova, p = 0.64
Anova, p = 0.0031
*
13.0
12.5
12.0
11.5
11.0
HEART CL(72:8)
Anova, p = 0.37
Anova, p = 2.8e-05
** **
12.5
12.0
11.5
11.0
D
SKMVL mito reads
Anova, p = 4.4e-06
* ** *** ***
Anova, p = 0.37
30
25
20
15
SKMVL mtDNA
Anova, p = 5.1e-05
**
Anova, p = 0.091
12.4
12.0
11.6
E
BAT mito reads
Anova, p = 0.013
*
Anova, p = 5.2e-06
***
50
40
30
20
BAT mtDNA
Anova, p = 0.039
*
Anova, p = 0.013
*
12
11
10
9

**Supplementary Figure 3. Training-induced changes in biomarkers of mitochondrial volume.** Visualization of biomarker data in **A)** Adrenal gland, **B)** Lung, **C)** Heart, **D)** Skeletal muscle, vastus lateralis (SKM-VL) and **E)** Brown adipose tissue (BAT). Biomarkers included are percent mitochondrial reads from RNA-seq (mito reads), mtDNA quantification from qRT-PCR analysis (mtDNA), and cardiolipin (CL) level for the most abundant species (72:8). CLs were available for selected tissues. Each boxplot represents the abundance level in a specific sex and time group. ANOVA statistics are provided for each tissue and sex combination. The whiskers extend from the hinge to the largest and lowest values, but no further than 1.5 * (the interquartile range).

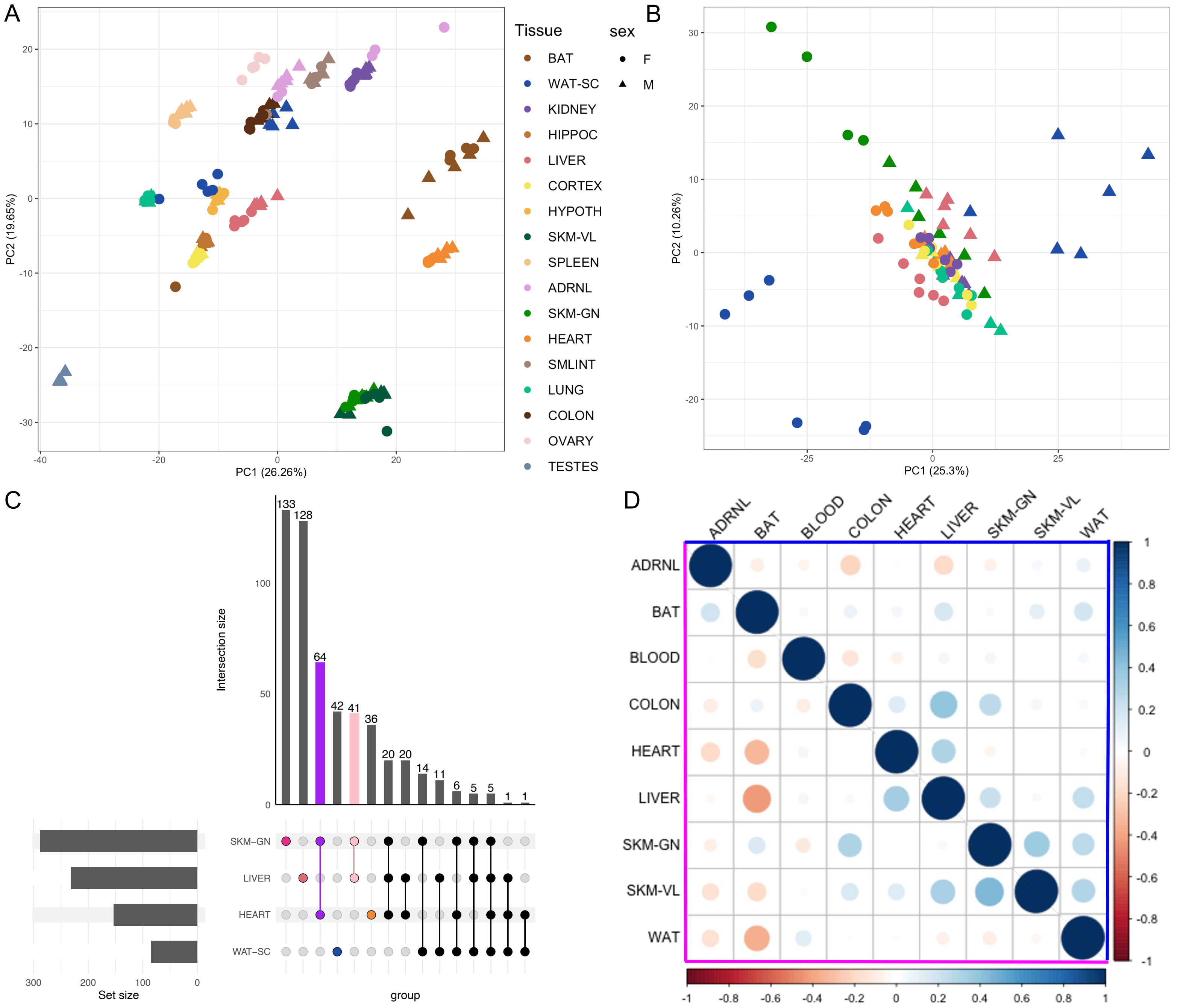
A
PC1 (26.26%)
PC2 (19.65%)
Tissue
BAT
WAT-SC
KIDNEY
HIPPOC
LIVER
CORTEX
HYPOTH
SKM-VL
SPLEEN
ADRNL
SKM-GN
HEART
SMLINT
LUNG
COLON
OVARY
TESTES
sex
F
M
B
PC1 (25.3%)
PC2 (10.26%)
C
Intersection size
133
128
64
42
41
36
20
20
14
11
6
5
5
1
1
SKM–GN
LIVER
HEART
WAT–SC
Set size
group
D
ADRNL
BAT
BLOOD
COLON
HEART
LIVER
SKM-GN
SKM-VL
WAT

**Supplementary Figure 4. The baseline and training-induced mitochondria-associated transcriptome and proteome. A-B)** Principal component analysis of the MitoCarta transcript (A) and protein (B) abundance for each tissue and animal in the untrained state. Samples are colored by tissue with females represented by circles and males by triangles. **C)** UpSet plot to characterize the number of training-differential MitoCarta proteins across the responding tissues that were analyzed for proteomics. Numbers above vertical bars indicate the number of proteins regulated by training in the tissues indicated by connected points below the bar. Horizontal bars indicate the total number of differential proteins in each tissue. **D)** Pearson correlation between gene expression changes (log2 fold change) for all MitoCarta genes after 8 weeks of endurance training in the 9 most responding tissues. Lower left corner shows the correlation for females (highlighted with a pink frame), and the upper right corner of the plot shows the correlation for males (highlighted with a blue frame).

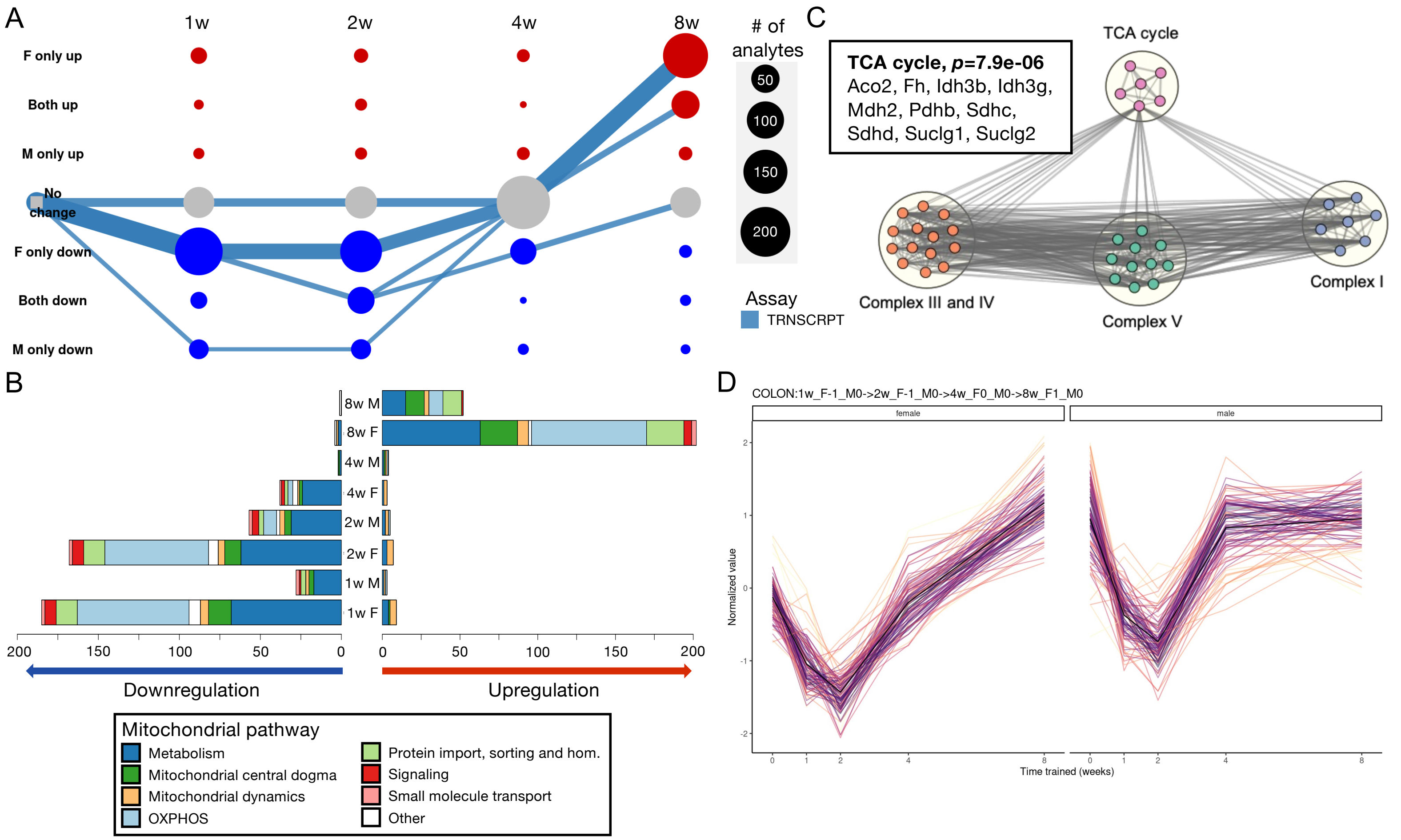
A
1w
2w
4w
8w
F only up
Both up
M only up
No change
F only down
Both down
M only down
### of analytes
50
100
150
200
Assay
TRNSCRPT
C
TCA cycle, p=7.9e-06
Aco2, Fh, Idh3b, Idh3g, Mdh2, Pdhb, Sdhc, Sdhd, Suclg1, Suclg2
TCA cycle
Complex III and IV
Complex V
Complex I
B
8w M
8w F
4w M
4w F
2w M
2w F
1w M
1w F
Downregulation
Upregulation
Mitochondrial pathway
Metabolism
Mitochondrial central dogma
Mitochondrial dynamics
OXPHOS
Protein import, sorting and hom.
Signaling
Small molecule transport
Other
D
COLON:1w_F-1_M0->2w_F-1_M0->4w_F0_M0->8w_F1_M0
female
male
Normalized value
Time trained (weeks)

**Supplementary figure 5. Training-induced changes in the colon. A)** Graphical representation of the top five mitochondria-associated training-differential trajectories in the colon. Each node represents one of nine possible states (row labels, with F for females and M for males, seven possible states are shown) at each of the four sampled training time points (column labels). Edges are drawn through these nodes to represent the path of differential analytes over the training time course, with edge color representing the –ome. The presented trajectories represent the top 5 clusters with greatest number of transcripts. Both node and edge size are proportional to the number of analytes represented by the node or edge. **B)** Number of significantly up- and downregulated mitochondria-associated colon transcripts at each training timepoint per sex, with color representation based on main MitoCarta pathway association of each transcript. **C)** Network view of pathway enrichment results corresponding to the largest differential path in the colon (1w_F-1_M0->2w_F-1_M0->4w_F0_M0->8w_F1_M0), representing downregulation in females during the early adaptation phase (1-2 weeks), through no change at 4 weeks and female-specific upregulation after 8 weeks of training). All enrichments are based on the transcriptome. Nodes indicate significantly enriched pathways (10% FDR), and an edge represents a pair of nodes with a similarity score of at least 0.3 between the gene sets driving each pathway enrichment. Clusters of enriched pathways were defined using Louvain community detection and are annotated with high-level biological themes. Node color indicates cluster assignment. Examples of genes in the TCA cycle pathway are highlighted within the framed square. **D)** Normalized expression levels of all analytes in largest differential path in the colon(1w_F-1_M0->2w_F-1_M0->4w_F0_M0->8w_F1_M0), representing downregulation in females during the early adaptation phase (1-2 weeks), through no change at 4 weeks and female-specific upregulation after 8 weeks of training).

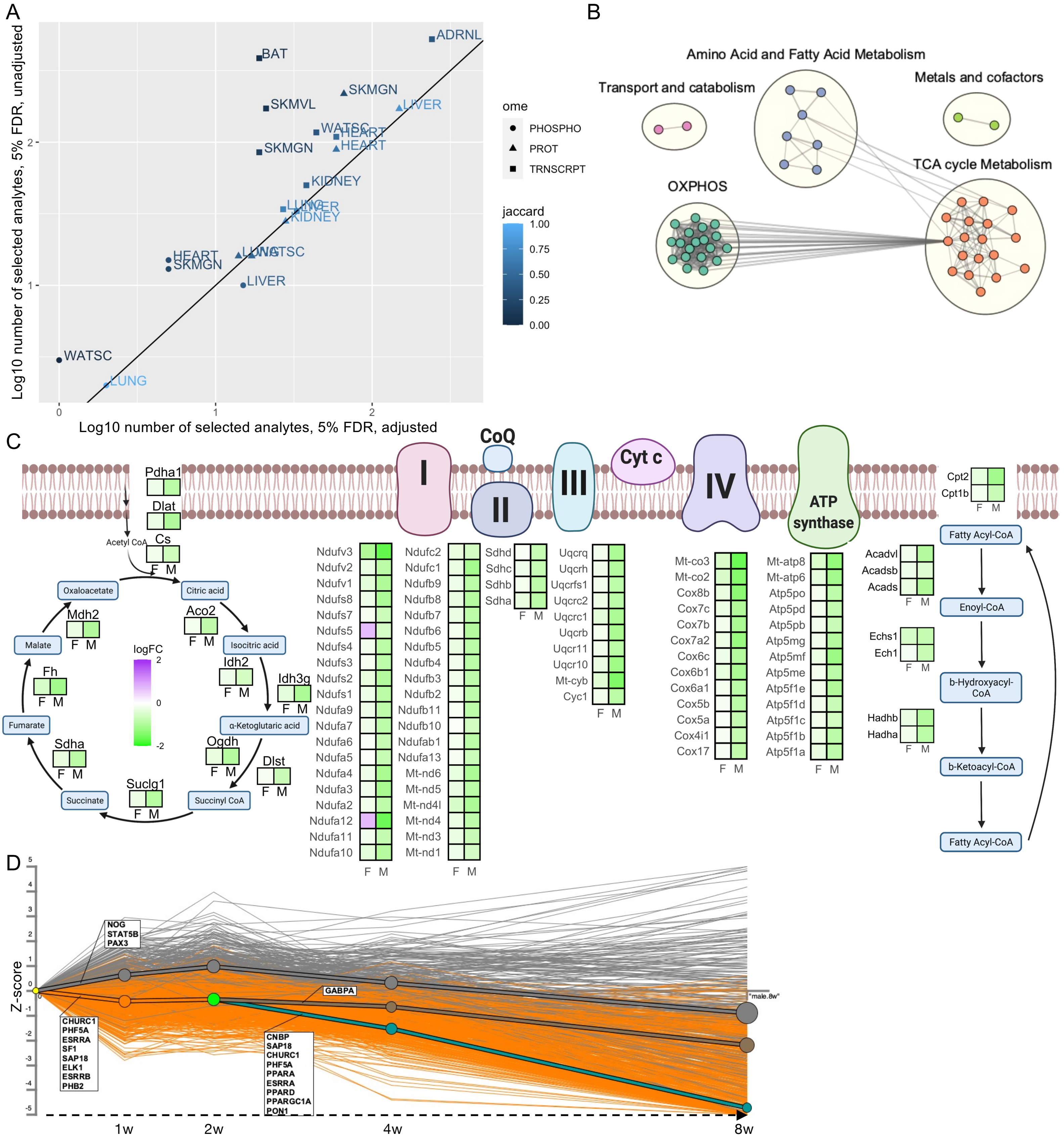

A
Log10 number of selected analytes, 5% FDR, unadjusted
Log10 number of selected analytes, 5% FDR, adjusted
ADRNL
BAT
SKMGN
SKMVL
LIVER
WATSC
HEART
SKMGN
KIDNEY
LUNG
LIVER
KIDNEY
HEART
SKMGN
LUNG
WATSC
LIVER
WATSC
LUNG
ome
PHOSPHO
PROT
TRNSCRPT
jaccard
1.00
0.75
0.50
0.25
0.00
B
Amino Acid and Fatty Acid Metabolism
Transport and catabolism
Metals and cofactors
TCA cycle Metabolism
OXPHOS
C
CoQ
Cyt c
I
II
III
IV
ATP synthase
Pdha1
Dlat
Acetyl CoA
Cs
Oxaloacetate
Citric acid
Mdh2
Aco2
Malate
Isocitric acid
Idh2
Idh3g
Fh
logFC
Fumarate
α-Ketoglutaric acid
Sdha
Ogdh
Dlst
Suclg1
Succinate
Succinyl CoA
F M
Ndufv3
Ndufv2
Ndufv1
Ndufs8
Ndufs7
Ndufs5
Ndufs4
Ndufs3
Ndufs2
Ndufs1
Ndufa9
Ndufa7
Ndufa6
Ndufa5
Ndufa4
Ndufa3
Ndufa2
Ndufa12
Ndufa11
Ndufa10
Ndufc2
Ndufc1
Ndufb9
Ndufb8
Ndufb7
Ndufb6
Ndufb5
Ndufb4
Ndufb3
Ndufb2
Ndufb11
Ndufb10
Ndufab1
Ndufa13
Mt-nd6
Mt-nd5
Mt-nd4l
Mt-nd4
Mt-nd3
Mt-nd1
Sdhd
Sdhc
Sdhb
Sdha
Uqcrq
Uqcrh
Uqcrfs1
Uqcrc2
Uqcrc1
Uqcrb
Uqcr11
Uqcr10
Mt-cyb
Cyc1
Mt-co3
Mt-co2
Cox8b
Cox7c
Cox7b
Cox7a2
Cox6c
Cox6b1
Cox6a1
Cox5b
Cox5a
Cox4i1
Cox17
Mt-atp8
Mt-atp6
Atp5po
Atp5pd
Atp5pb
Atp5mg
Atp5mf
Atp5me
Atp5f1e
Atp5f1d
Atp5f1c
Atp5f1b
Atp5f1a
Cpt2
Cpt1b
Fatty Acyl-CoA
Acadvl
Acadsb
Acads
Enoyl-CoA
Echs1
Ech1
b-Hydroxyacyl-CoA
Hadhb
Hadha
b-Ketoacyl-CoA
Fatty Acyl-CoA
D
Z-score
NOG
STAT5B
PAX3
CHURC1
PHF5A
ESRRA
SF1
SAP18
ELK1
ESRRB
PHB2
GABPA
CNBP
SAP18
CHURC1
PHF5A
PPARA
ESRRA
PPARD
PPARGC1A
PON1
"male.8w"
1w
2w
4w
8w

**Supplementary Figure 6. Mitochondrial content partially explains response in BAT and the adrenal gland. A)** The (log10) number of mitochondria-associated multi-omic analytes selected at 5% FDR control, comparing the unadjusted (y-axis) to the biomarker-adjusted (x-axis) analyses. **B)** Network view of pathway enrichment results corresponding to the largest differential path in the BAT (1w_F0_M0->2w_F0_M0->4w_F0_M0->8w_F-1_M-1), representing no change up to 4 weeks of training, followed by a late downregulation of mitochondria-associated analytes after 8 weeks of training. Nodes indicate significantly enriched pathways (10% FDR), and an edge represents a pair of nodes with a similarity score of at least 0.3 between the gene sets driving each pathway enrichment. Clusters of enriched pathways were defined using Louvain community detection and are annotated with high-level biological themes. Node color indicates cluster assignment. **C)** Individual transcript changes (log2 fold change) for the main mitochondrial ATP-producing pathways (beta-oxidation, TCA cycle and electron transport chain). All transcripts associated with these pathways that were significantly regulated in males after 8 weeks of training are shown, with color scale indicating degree of change. **D)** Dynamic regulatory events miner (DREM) analysis results of the MitoCarta genes in male BAT predict several transcription factors to be involved in the late downregulation that occurs from 2 to 8 weeks of training. Y-axis represents timewise z-scores extracted from our unadjusted differential analysis results.

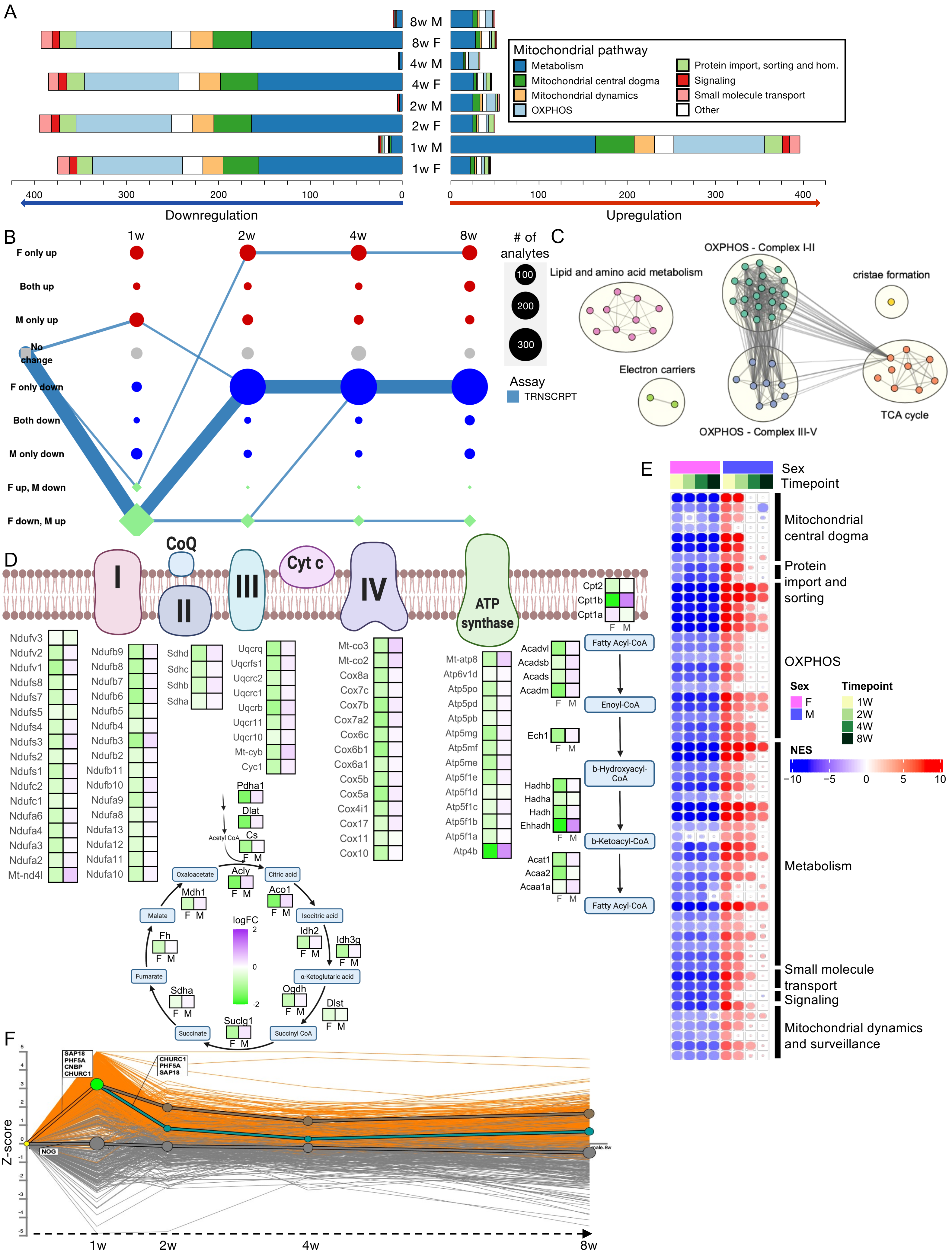
A
8w M
8w F
4w M
4w F
2w M
2w F
1w M
1w F
Mitochondrial pathway
Metabolism
Mitochondrial central dogma
Mitochondrial dynamics
OXPHOS
Protein import, sorting and hom.
Signaling
Small molecule transport
Other
400
300
200
100
0
Downregulation
Upregulation
B
1w
2w
4w
8w
F only up
Both up
M only up
No change
F only down
Both down
M only down
F up, M down
F down, M up
### of analytes
100
200
300
Assay
TRNSCRPT
C
Lipid and amino acid metabolism
OXPHOS - Complex I-II
cristae formation
Electron carriers
TCA cycle
OXPHOS - Complex III-V
D
CoQ
Cyt c
I
II
III
IV
ATP synthase
Cpt2
Cpt1b
Cpt1a
Fatty Acyl-CoA
Enoyl-CoA
b-Hydroxyacyl-CoA
b-Ketoacyl-CoA
Fatty Acyl-CoA
Acadvl
Acadsb
Acads
Acadm
Ech1
Hadhb
Hadha
Hadh
Ehhadh
Acat1
Acaa2
Acaa1a
F M
Ndufv3
Ndufv2
Ndufv1
Ndufs8
Ndufs7
Ndufs5
Ndufs4
Ndufs3
Ndufs2
Ndufs1
Ndufc2
Ndufc1
Ndufa6
Ndufa4
Ndufa3
Ndufa2
Mt-nd4l
Ndufb9
Ndufb8
Ndufb7
Ndufb6
Ndufb5
Ndufb4
Ndufb3
Ndufb2
Ndufb11
Ndufb10
Ndufa9
Ndufa8
Ndufa13
Ndufa12
Ndufa11
Ndufa10
Sdhd
Sdhc
Sdhb
Sdha
Uqcrq
Uqcrfs1
Uqcrc2
Uqcrc1
Uqcrb
Uqcr11
Uqcr10
Mt-cyb
Cyc1
Mt-co3
Mt-co2
Cox8a
Cox7c
Cox7b
Cox7a2
Cox6c
Cox6b1
Cox6a1
Cox5b
Cox5a
Cox4i1
Cox17
Cox11
Cox10
Mt-atp8
Atp6v1d
Atp5po
Atp5pd
Atp5pb
Atp5mg
Atp5mf
Atp5me
Atp5f1e
Atp5f1d
Atp5f1c
Atp5f1b
Atp5f1a
Atp4b
Pdha1
Dlat
Acetyl CoA
Cs
Oxaloacetate
Acly
Citric acid
Mdh1
Aco1
Malate
Isocitric acid
logFC
2
0
-2
Idh2
Idh3g
Fh
Fumarate
α-Ketoglutaric acid
Sdha
Ogdh
Dlst
Succinate
Suclg1
Succinyl CoA
E
Sex
Timepoint
Mitochondrial central dogma
Protein import and sorting
OXPHOS
Sex
F
M
Timepoint
1W
2W
4W
8W
NES
−10
−5
0
5
10
Metabolism
Small molecule transport
Signaling
Mitochondrial dynamics and surveillance
F
SAP18
PHF5A
CNBP
CHURC1
CHURC1
PHF5A
SAP18
NOG
Z-score
1w
2w
4w
8w

**Supplementary figure 7. Training remodels the mitochondria-associated transcriptome in the adrenal gland. A)** Number of significantly up- and down-regulated mitochondria-associated adrenal transcripts at each training timepoint per sex, with color representation based on main MitoCarta pathway association of each transcript. **B)** Graphical representation of the mitochondria-associated training-differential analytes in the adrenal glands. Each node represents one of nine possible states (row labels, with F for females and M for males) at each of the four sampled training time points (column labels). Edges are drawn through these nodes to represent the path of differential analytes over the training time course, with edge color representing the –ome. This graph includes the five largest paths for the adrenal glands. Both node and edge size are proportional to the number of analytes represented by the node or edge. **C)** Network view of pathway enrichment results corresponding to the largest differential path in the adrenal glands (1w_F-1_M1->2w_F-1_M0->4w_F-1_M0->8w_F-1_M0), representing downregulation in females across the 8-week timecourse, and upregulation in males after 1 week before going back to baseline. All enrichments are based on the transcriptomics results. Nodes indicate significantly enriched pathways (10% FDR), and an edge represents a pair of nodes with a similarity score of at least 0.3 between the gene sets driving each pathway enrichment. Clusters of enriched pathways were defined using Louvain community detection and are annotated with high-level biological themes. Node color indicates cluster assignment. **D)** Individual transcript changes (log2 fold change) for the main mitochondrial ATP-producing pathways (beta-oxidation, TCA cycle and electron transport chain). All transcripts associated with these pathways that were significantly regulated in females after 8 weeks of training are shown, with color scale indicating degree of change. **E)** Gene Set Enrichment Analysis (GSEA) results from the adrenal gland transcriptome dataset using the MitoCarta Mitopathways 3.0 gene set database. Individual pathways (rows) are organized and labelled based on main MitoCarta pathway category. Pathways included are significant (FDR<0.01) in both sexes after 1 week of training. Heatmaps show NES as color for each timepoint for both sexes. **F)** Dynamic regulatory events miner (DREM) analysis results of the MitoCarta genes in male adrenal gland predict several transcription factors to be involved in the early (1w) responses. Y-axis represents timewise z-scores extracted from our unadjusted differential analysis results.

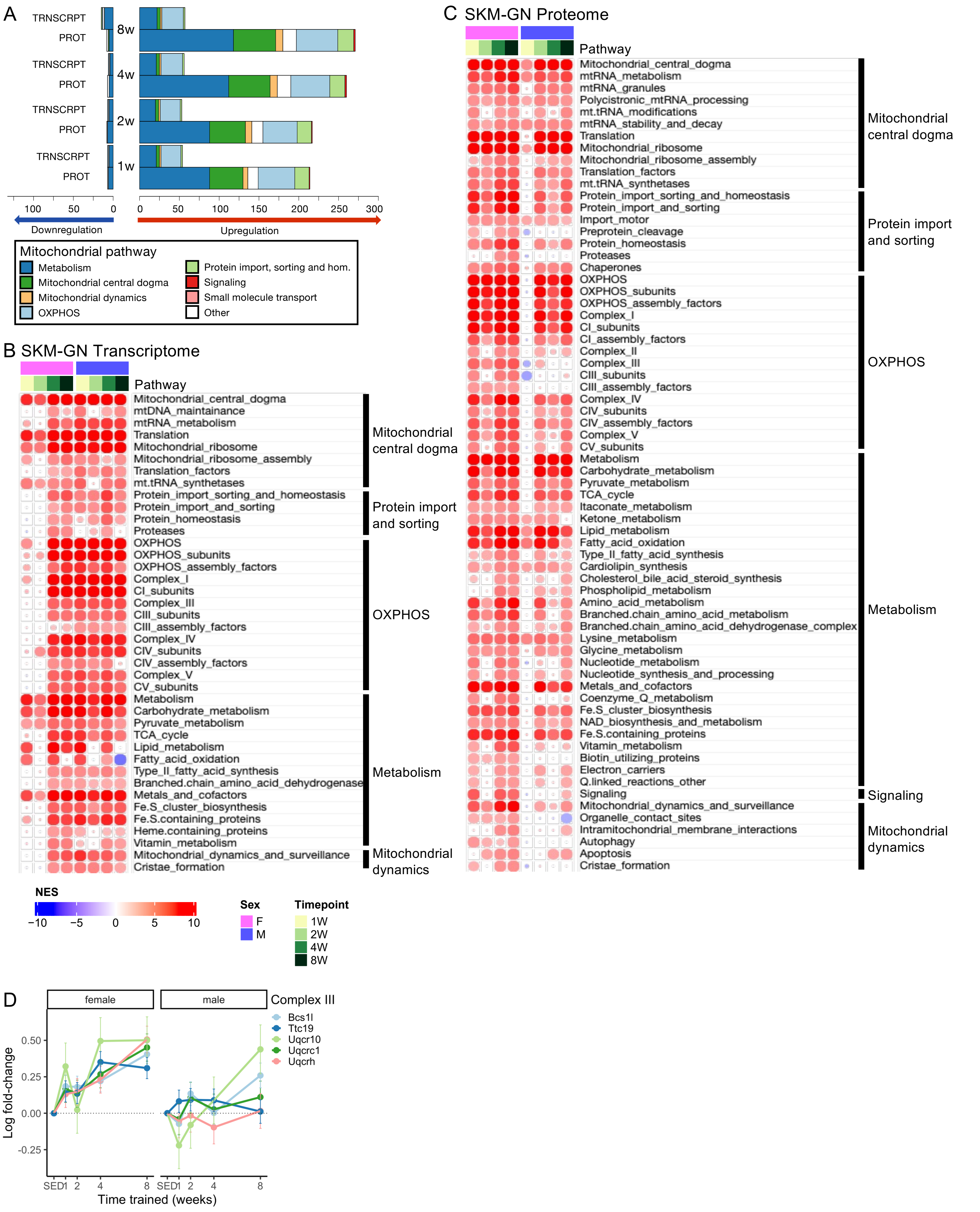

A
TRNSCRPT
PROT
8w
4w
2w
1w
100 50 0
0 50 100 150 200 250 300
Downregulation
Upregulation
Mitochondrial pathway
Metabolism
Mitochondrial central dogma
Mitochondrial dynamics
OXPHOS
Protein import, sorting and hom.
Signaling
Small molecule transport
Other
B SKM-GN Transcriptome
Pathway
Mitochondrial_central_dogma
mtDNA_maintainance
mtRNA_metabolism
Translation
Mitochondrial_ribosome
Mitochondrial_ribosome_assembly
Translation_factors
mt.tRNA_synthetases
Protein_import_sorting_and_homeostasis
Protein_import_and_sorting
Protein_homeostasis
Proteases
OXPHOS
OXPHOS_subunits
OXPHOS_assembly_factors
Complex_I
CI_subunits
Complex_III
CIII_subunits
CIII_assembly_factors
Complex_IV
CIV_subunits
CIV_assembly_factors
Complex_V
CV_subunits
Metabolism
Carbohydrate_metabolism
Pyruvate_metabolism
TCA_cycle
Lipid_metabolism
Fatty_acid_oxidation
Type_II_fatty_acid_synthesis
Branched.chain_amino_acid_dehydrogenase
Metals_and_cofactors
Fe.S_cluster_biosynthesis
Fe.S.containing_proteins
Heme.containing_proteins
Vitamin_metabolism
Mitochondrial_dynamics_and_surveillance
Cristae_formation
Mitochondrial central dogma
Protein import and sorting
OXPHOS
Metabolism
Mitochondrial dynamics
C SKM-GN Proteome
Pathway
Mitochondrial_central_dogma
mtRNA_metabolism
mtRNA_granules
Polycistronic_mtRNA_processing
mt.tRNA_modifications
mtRNA_stability_and_decay
Translation
Mitochondrial_ribosome
Mitochondrial_ribosome_assembly
Translation_factors
mt.tRNA_synthetases
Protein_import_sorting_and_homeostasis
Protein_import_and_sorting
Import_motor
Preprotein_cleavage
Protein_homeostasis
Proteases
Chaperones
OXPHOS
OXPHOS_subunits
OXPHOS_assembly_factors
Complex_I
CI_subunits
CI_assembly_factors
Complex_II
Complex_III
CIII_subunits
CIII_assembly_factors
Complex_IV
CIV_subunits
CIV_assembly_factors
Complex_V
CV_subunits
Metabolism
Carbohydrate_metabolism
Pyruvate_metabolism
TCA_cycle
Itaconate_metabolism
Ketone_metabolism
Lipid_metabolism
Fatty_acid_oxidation
Type_II_fatty_acid_synthesis
Cardiolipin_synthesis
Cholesterol_bile_acid_steroid_synthesis
Phospholipid_metabolism
Amino_acid_metabolism
Branched.chain_amino_acid_metabolism
Branched.chain_amino_acid_dehydrogenase_complex
Lysine_metabolism
Glycine_metabolism
Nucleotide_metabolism
Nucleotide_synthesis_and_processing
Metals_and_cofactors
Coenzyme_Q_metabolism
Fe.S_cluster_biosynthesis
NAD_biosynthesis_and_metabolism
Fe.S.containing_proteins
Vitamin_metabolism
Biotin_utilizing_proteins
Electron_carriers
Q.linked_reactions_other
Signaling
Mitochondrial_dynamics_and_surveillance
Organelle_contact_sites
Intramitochondrial_membrane_interactions
Autophagy
Apoptosis
Cristae_formation
Mitochondrial central dogma
Protein import and sorting
OXPHOS
Metabolism
Signaling
Mitochondrial dynamics
NES
−10 −5 0 5 10
Sex
F
M
Timepoint
1W
2W
4W
8W
D
female
male
Complex III
Bcs1l
Ttc19
Uqcr10
Uqcrc1
Uqcrh
Log fold-change
0.50
0.25
0.00
-0.25
SED 1 2 4 8
Time trained (weeks)

**Supplementary figure 8. Concordant mitochondrial response in male and female skeletal muscle. A)** Number of significantly up- and downregulated analytes at each training timepoint, with color representation based on main MitoCarta pathway association of each analyte, in SKM-GN transcripts and proteins. **B-C)** Gene Set Enrichment Analysis (GSEA) results from the gastrocnemius (SKM-GN) transcriptome (B), and global proteome (C) datasets using the MitoCarta Mitopathways 3.0 gene set database. Individual pathways (rows) are organized and labelled based on main MitoCarta pathway category. Pathways included are significant (FDR<0.001) in at least one sex after 8 weeks of training. Heatmaps show normalized enrichment scores (NES) as color for each timepoint for both sexes. **D)** Log2 fold change of protein expression changes in Complex III proteins, driving the female-specific pathway enrichment. All proteins are significant in females, whereas only Bcs1l and Uqcr10 are significant in males after 8 weeks.

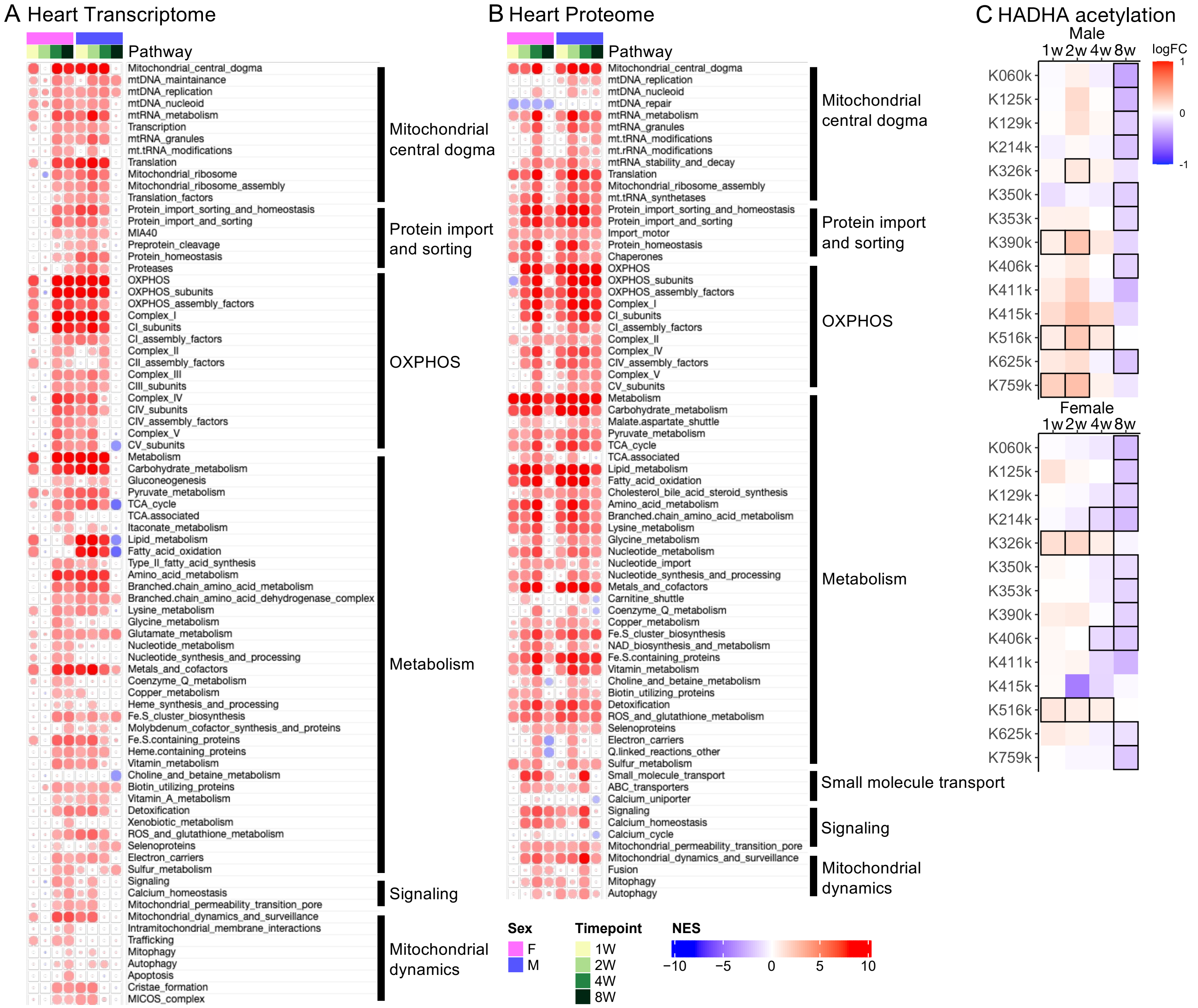

A Heart Transcriptome
Pathway
Mitochondrial_central_dogma
mtDNA_maintainance
mtDNA_replication
mtDNA_nucleoid
mtRNA_metabolism
Transcription
mtRNA_granules
mt.tRNA_modifications
Translation
Mitochondrial_ribosome
Mitochondrial_ribosome_assembly
Translation_factors
Protein_import_sorting_and_homeostasis
Protein_import_and_sorting
MIA40
Preprotein_cleavage
Protein_homeostasis
Proteases
OXPHOS
OXPHOS_subunits
OXPHOS_assembly_factors
Complex_I
CI_subunits
CI_assembly_factors
Complex_II
CII_assembly_factors
Complex_III
CIII_subunits
Complex_IV
CIV_subunits
CIV_assembly_factors
Complex_V
CV_subunits
Metabolism
Carbohydrate_metabolism
Gluconeogenesis
Pyruvate_metabolism
TCA_cycle
TCA.associated
Itaconate_metabolism
Lipid_metabolism
Fatty_acid_oxidation
Type_II_fatty_acid_synthesis
Amino_acid_metabolism
Branched.chain_amino_acid_metabolism
Branched.chain_amino_acid_dehydrogenase_complex
Lysine_metabolism
Glycine_metabolism
Glutamate_metabolism
Nucleotide_metabolism
Nucleotide_synthesis_and_processing
Metals_and_cofactors
Coenzyme_Q_metabolism
Copper_metabolism
Heme_synthesis_and_processing
Fe.S_cluster_biosynthesis
Molybdenum_cofactor_synthesis_and_proteins
Fe.S.containing_proteins
Heme.containing_proteins
Vitamin_metabolism
Choline_and_betaine_metabolism
Biotin_utilizing_proteins
Vitamin_A_metabolism
Detoxification
Xenobiotic_metabolism
ROS_and_glutathione_metabolism
Selenoproteins
Electron_carriers
Sulfur_metabolism
Signaling
Calcium_homeostasis
Mitochondrial_permeability_transition_pore
Mitochondrial_dynamics_and_surveillance
Intramitochondrial_membrane_interactions
Trafficking
Mitophagy
Autophagy
Apoptosis
Cristae_formation
MICOS_complex
Mitochondrial central dogma
Protein import and sorting
OXPHOS
Metabolism
Signaling
Mitochondrial dynamics
B Heart Proteome
Pathway
Mitochondrial_central_dogma
mtDNA_replication
mtDNA_nucleoid
mtDNA_repair
mtRNA_metabolism
mtRNA_granules
mt.tRNA_modifications
mt.rRNA_modifications
mtRNA_stability_and_decay
Translation
Mitochondrial_ribosome_assembly
mt.tRNA_synthetases
Protein_import_sorting_and_homeostasis
Protein_import_and_sorting
Import_motor
Protein_homeostasis
Chaperones
OXPHOS
OXPHOS_subunits
OXPHOS_assembly_factors
Complex_I
CI_subunits
CI_assembly_factors
Complex_II
Complex_IV
CIV_assembly_factors
Complex_V
CV_subunits
Metabolism
Carbohydrate_metabolism
Malate.aspartate_shuttle
Pyruvate_metabolism
TCA_cycle
TCA.associated
Lipid_metabolism
Fatty_acid_oxidation
Cholesterol_bile_acid_steroid_synthesis
Amino_acid_metabolism
Branched.chain_amino_acid_metabolism
Lysine_metabolism
Glycine_metabolism
Nucleotide_metabolism
Nucleotide_import
Nucleotide_synthesis_and_processing
Metals_and_cofactors
Carnitine_shuttle
Coenzyme_Q_metabolism
Copper_metabolism
Fe.S_cluster_biosynthesis
NAD_biosynthesis_and_metabolism
Fe.S.containing_proteins
Vitamin_metabolism
Choline_and_betaine_metabolism
Biotin_utilizing_proteins
Detoxification
ROS_and_glutathione_metabolism
Selenoproteins
Electron_carriers
Q.linked_reactions_other
Sulfur_metabolism
Small_molecule_transport
ABC_transporters
Calcium_uniporter
Signaling
Calcium_homeostasis
Calcium_cycle
Mitochondrial_permeability_transition_pore
Mitochondrial_dynamics_and_surveillance
Fusion
Mitophagy
Autophagy
Mitochondrial central dogma
Protein import and sorting
OXPHOS
Metabolism
Small molecule transport
Signaling
Mitochondrial dynamics
Sex
F
M
Timepoint
1W
2W
4W
8W
NES
−10
−5
0
5
10
C HADHA acetylation
Male
1w 2w 4w 8w
K060k
K125k
K129k
K214k
K326k
K350k
K353k
K390k
K406k
K411k
K415k
K516k
K625k
K759k
Female
1w 2w 4w 8w
K060k
K125k
K129k
K214k
K326k
K350k
K353k
K390k
K406k
K411k
K415k
K516k
K625k
K759k
logFC
1
0
-1

**Supplementary Figure 9. Mitochondrial adaptation in the heart muscle.** Gene Set Enrichment Analysis (GSEA) results from **A)** the heart transcriptome, and **B)** the heart global proteome datasets using the MitoCarta Mitopathways 3.0 gene set database. Individual pathways (rows) are organized and labelled based on main MitoCarta pathway category. Pathways included are significant (FDR<0.05) in either sex (or both) after 8 weeks of training. Heatmaps show normalized enrichment scores (NES) as color for each timepoint for both sexes. **C)** Site-specific acetylation changes in HADHA in males (top panel) and females (bottom panel). All displayed sites were differentially acetylated overall (taking all timepoints and sexes into account, FDR<0.05), and sites that reach timewise significance (FDR<0.05) are highlighted with black frames.

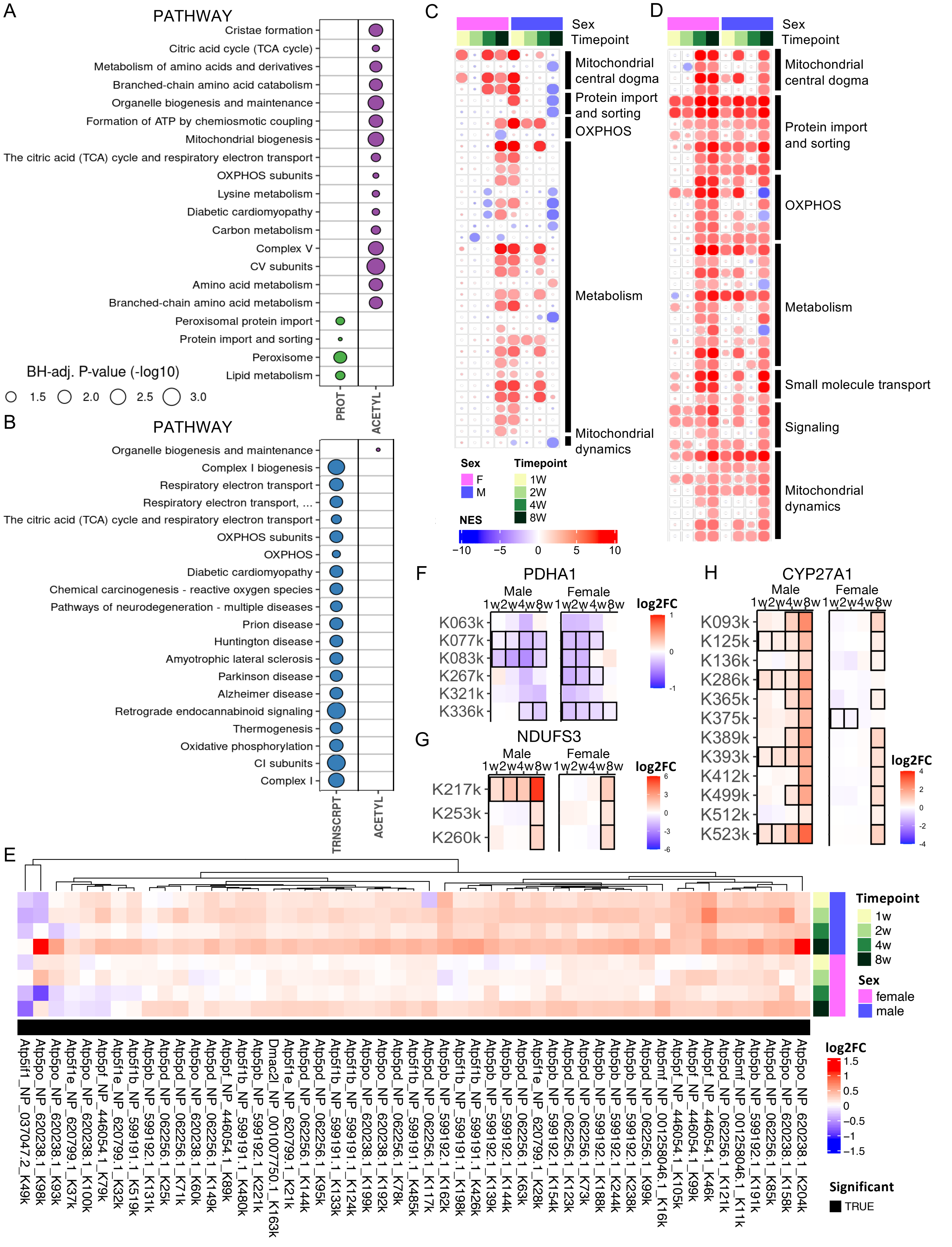
A
PATHWAY
Cristae formation
Citric acid cycle (TCA cycle)
Metabolism of amino acids and derivatives
Branched-chain amino acid catabolism
Organelle biogenesis and maintenance
Formation of ATP by chemiosmotic coupling
Mitochondrial biogenesis
The citric acid (TCA) cycle and respiratory electron transport
OXPHOS subunits
Lysine metabolism
Diabetic cardiomyopathy
Carbon metabolism
Complex V
CV subunits
Amino acid metabolism
Branched-chain amino acid metabolism
Peroxisomal protein import
Protein import and sorting
Peroxisome
Lipid metabolism
BH-adj. P-value (-log10)
1.5
2.0
2.5
3.0
PROT
ACETYL
B
PATHWAY
Organelle biogenesis and maintenance
Complex I biogenesis
Respiratory electron transport
Respiratory electron transport, ...
The citric acid (TCA) cycle and respiratory electron transport
OXPHOS subunits
OXPHOS
Diabetic cardiomyopathy
Chemical carcinogenesis - reactive oxygen species
Pathways of neurodegeneration - multiple diseases
Prion disease
Huntington disease
Amyotrophic lateral sclerosis
Parkinson disease
Alzheimer disease
Retrograde endocannabinoid signaling
Thermogenesis
Oxidative phosphorylation
CI subunits
Complex I
TRNSCRPT
ACETYL
C
Sex
Timepoint
Mitochondrial central dogma
Protein import and sorting
OXPHOS
Metabolism
Mitochondrial dynamics
Sex
F
M
Timepoint
1W
2W
4W
8W
NES
−10
−5
0
5
10
D
Sex
Timepoint
Mitochondrial central dogma
Protein import and sorting
OXPHOS
Metabolism
Small molecule transport
Signaling
Mitochondrial dynamics
F
PDHA1
Male
Female
1w2w4w8w1w2w4w8w
K063k
K077k
K083k
K267k
K321k
K336k
log2FC
1
0
-1
G
NDUFS3
Male
Female
1w2w4w8w1w2w4w8w
K217k
K253k
K260k
log2FC
6
3
0
-3
-6
H
CYP27A1
Male
Female
1w2w4w8w1w2w4w8w
K093k
K125k
K136k
K286k
K365k
K375k
K389k
K393k
K412k
K499k
K512k
K523k
log2FC
4
2
0
-2
-4
E
Timepoint
1w
2w
4w
8w
Sex
female
male
Atp5if1_NP_037047.2_K49k
Atp5po_NP_620238.1_K98k
Atp5po_NP_620238.1_K93k
Atp5f1e_NP_620799.1_K37k
Atp5po_NP_620238.1_K100k
Atp5pf_NP_446054.1_K79k
Atp5f1e_NP_620799.1_K32k
Atp5f1b_NP_599191.1_K519k
Atp5pb_NP_599192.1_K131k
Atp5pd_NP_062256.1_K25k
Atp5pd_NP_062256.1_K71k
Atp5po_NP_620238.1_K60k
Atp5pd_NP_062256.1_K149k
Atp5pf_NP_446054.1_K89k
Atp5f1b_NP_599191.1_K480k
Atp5pb_NP_599192.1_K221k
Dmac2l_NP_001007750.1_K163k
Atp5f1e_NP_620799.1_K21k
Atp5pd_NP_062256.1_K144k
Atp5pd_NP_062256.1_K95k
Atp5f1b_NP_599191.1_K133k
Atp5f1b_NP_599191.1_K124k
Atp5po_NP_620238.1_K199k
Atp5po_NP_620238.1_K192k
Atp5pd_NP_062256.1_K78k
Atp5f1b_NP_599191.1_K485k
Atp5pd_NP_062256.1_K117k
Atp5pb_NP_599192.1_K162k
Atp5f1b_NP_599191.1_K198k
Atp5f1b_NP_599191.1_K426k
Atp5pb_NP_599192.1_K139k
Atp5pb_NP_599192.1_K144k
Atp5pd_NP_062256.1_K63k
Atp5f1e_NP_620799.1_K28k
Atp5pb_NP_599192.1_K154k
Atp5pd_NP_062256.1_K123k
Atp5pd_NP_062256.1_K73k
Atp5pb_NP_599192.1_K188k
Atp5pb_NP_599192.1_K244k
Atp5pb_NP_599192.1_K238k
Atp5pd_NP_062256.1_K99k
Atp5mf_NP_001258046.1_K16k
Atp5pf_NP_446054.1_K105k
Atp5pf_NP_446054.1_K99k
Atp5pf_NP_446054.1_K46k
Atp5pd_NP_062256.1_K121k
Atp5mf_NP_001258046.1_K11k
Atp5pb_NP_599192.1_K191k
Atp5pd_NP_062256.1_K85k
Atp5po_NP_620238.1_K158k
Atp5po_NP_620238.1_K204k
log2FC
1.5
1
0.5
0
−0.5
−1
−1.5
Significant
TRUE

**Supplementary Figure 10. Liver.** Pathway enrichment analysis of **A)** the largest differential path (1w_F0_M1->2w_F0_M1->4w_F0_M1->8w_F1_M1, representing upregulation in males only for the 1, 2 and 4 week timepoints, followed by an increase in both sexes after 8 weeks) and **B)** the second largest path (1w_F0_M0->2w_F0_M0->4w_F0_M0->8w_F1_M1, represeting no change during the first 4 weeks of training, followed by an increase in both sexes after 8 weeks) in the liver. If no pathways are shown for a specific –ome, that means no significant enrichments were identified. Node size is proportional to the BH-adjusted p-value of the enrichment. **C)** Gene Set Enrichment Analysis (GSEA) results from the liver transcriptome dataset using the MitoCarta Mitopathways 3.0 gene set database. Individual pathways (rows) are organized and labelled based on main MitoCarta pathway category. Pathways included are significant (FDR<0.05) in either sex after 8 weeks of training. Heatmap shows normalized enrichment scores (NES) as color for each timepoint for both sexes. **D)** GSEA results from the liver proteome dataset using the MitoCarta Mitopathways 3.0 gene set database. Individual pathways (rows) are organized and labelled based on main MitoCarta pathway category. Pathways included are significant (FDR<0.01) in both sexes after 8 weeks of training. The more stringent cutoff compared to the transcriptome is used to limit the number of pathways for visualization purposes. Heatmap shows NES as color for each timepoint for both sexes. **E)** Complex V subunit pathway enrichment for the liver acetylome. Only significant acetylsites are shown. Rows are clustered using hierarchical clustering. Heatmap shows log2 fold change for each timepoint and sex. **F-H)** Site-specific acetylation changes in **F)** CYP27A1, **G)** NDUFS3 and **H)** PDHA1 in males and females. All displayed sites were differentially acetylated overall (taking all timepoints and sexes into account, FDR<0.05), and sites that reach timewise significance (FDR<0.05) are highlighted with black frames.
