## Supplementary text for "The mitochondrial multi-omic response to exercise training across tissues"

### Data generation and processing

#### RNA sequencing

Following tissue lysis, total RNA was extracted in a BiomekFx automation workstation. Total RNA from blood was extracted using the Agencourt RNAdvance blood specific kit (Beckman Coulter). The RNA quantity and quality were assessed with NanoDrop (ThermoFisher Scientific, # ND-ONE-W), Qubit assay (ThermoFisher Scientific), and either Bioanalyzer or Fragment Analyzer. Approximately 500 ng of total RNA was used to generate RNA-seq libraries from poly(A)-selected RNA using the Universal Plus mRNA-Seq kit from NuGEN/Tecan (# 9133). All libraries were prepared using a Biomek i7 laboratory automation system (Beckman Coulter). Pooled libraries were subsequently sequenced using 100bp paired-end sequencing on an Illumina NovaSeq 6000 platform (Illumina, San Diego, CA, USA) targeting a depth of 35 million read pairs per sample. Reads were demultiplexed with bcl2fastq2 (v2.20.0), adapters were trimmed with cutadapt (v1.18), pre-alignment QC metrics generated with FastQC (v0.11.8) and reads aligned using STAR (v2.7.0d). Quantification was performed using RSEM (v1.3.1). Normalized sample-level data were generated using filtered raw counts that were TMM-normalized, and converted to log counts per million with *edgeR* [1].

#### Reduced representation bisulfite sequencing (RRBS)

Tissues were lysed and DNA extracted using a BiomekFx automation workstation (Beckman Coulter). Approximately 100ng of DNA was used for library preparation using the Ovation® RRBS Methyl-Seq kit (Tecan Genomics), which includes bisulfite conversion of the DNA. The quantity and quality of the libraries were assessed using Qubit High Sensitivity assays (ThermoFisher Scientific), and the Bioanalyzer High Sensitivity DNA Chip (Agilent Technologies). The pooled libraries were sequenced using 100bp paired-end sequencing on an Illumina NovaSeq 6000 platform (Illumina, San Diego, CA, USA) to reach at least 30 million paired-end reads. Processing included demultiplexing with bcl2fastq (version 2.20), adapter trimming using TrimGalore (v1.18), and indexing and aligning reads to the *Rattus norvegicus* (rn6) genome using Bismark (v0.20.0). Normalization was performed using edgeR [1] sample-level data were generated from $log_2$-transformed filtered sites/clusters that were then quantile normalized using PreprocessCore.

#### Assay for transposase-accessible chromatin using sequencing (ATAC-seq)

Nuclei from aliquoted tissue samples were extracted using the Omni-ATAC protocol with modifications [2]. For all tissues, the homogenate passed through a 40 µm cell strainer to collect nuclei, that were then stained with DAPI and counted. 50,000 nuclei (or max. 500 µl nuclei) was mixed with 1 ml ATAC-RSB buffer and spun at 1000x*g* for 10 minutes. The nuclei pellet was resuspended in 50 µl of transposition mixture and incubated at 37°C for 30 minutes with 1000 rpm shaking. The transposed DNA was subsequently purified using the Qiagen MinElute

Purification kit (Qiagen # 28006). The DNA product was amplified using custom indexed primers and the NEBnext High-Fidelity 2x PCR Master Mix (NEB, M0541L), and cleaned with the 1.8x SPRIselect beads prior to sequencing. Pooled libraries were sequenced using 50bp paired-end sequencing on an Illumina NovaSeq 6000 platform (Illumina, San Diego, CA, USA) targeting a depth of 35 million read pairs per sample.Reads were demultiplexed with bcl2fastq2 (v2.20.0) and processed with the ENCODE ATAC-seq pipeline (v1.7.0) (https://github.com/ENCODE-DCC/atac-seq-pipeline) [3]. Adapter-trimming was performed using cutadapt (v2.5), and reads were aligned using Bowtie 2 (v2.3.4.3). Duplicate reads and reads mapping to the mitochondrial chromosome were removed. Signal files and peak calls were generated using MACS2 (v2.2.4).

### Proteomics

The BCA assay (ThermoFisher) was used to determine protein lysate concentrations. After reduction in 5 mM dithiothreitol (DTT, Sigma-Aldrich) for 1 hour at 37 °C under 1000 rpm mixing, the lysate was alkylated with iodoacetamide (IAA, Sigma-Aldrich) in the dark for 45 minutes at 25 °C under 1000 rpm mixing. All samples were diluted 1:4 with 50 mM Tris-HCl, pH 8.0 and LysC endopeptidase (1mAU/uL, Wako Chemicals) was added at a 1:50 enzyme:substrate ratio, followed by 2 hours of digestion at 25°C and 850 rpm mixing. The samples were then trypsinized at an enzyme:substrate ratio of 1:50 (1:10 for white adipose tissue) for 14 hours at 25 °C and 850 rpm mixing. Digestion was terminated by the addition of formic acid (FA) to reach a 1% total concentration, followed by centrifugation for 15 min at 1500x*g* at 4 °C. The resulting supernatant was first diluted with 0.1% FA to 3 ml total volume and desalted using Sep-Pac C18 SPE cartridges (Waters), and then concentrated using a speedvac prior to a new final concentration measurement through BCA assay.

Protein samples (400 µg) were first dried, and then resuspended in 200 mM HEPES pH 8.5 (final concentration 5 µg/uL) for TMT labeling. Samples were randomized into ten TMT channels, with the last channel (total 11 plexes) containing a common reference aliquot consisting of pooled peptides from each experimental sample. The TMT reagents and peptide aliquots were combined to reach a 1:1 peptide:tag ratio and mixed for 1 hour at 2 5°C, 400 rpm. Samples were then diluted to 2.5 µg/uL (in 20% acetonitrile). After QC checks, reactions were quenched with hydroxylamine and samples from each multiplex were combined, concentrated in a speedvac, and desalted using Sep-Pac C18 SPE cartridges (Waters). Heart and liver tissues were also subjected to phosphotyrosine enrichment as previously described [4]. Additional details on the processing have been published elsewhere[5].

After fractionation by high pH reversed phase separation, samples were resuspended in mobile phase A, centrifuged to remove any debris, and then loaded onto a column. The flow rate through the column was set to 1 ml/min (using mobile phase B), and samples were eluted for 96 minutes, resulting in the collection of 96 fractions. Global proteome analysis was run on 5 % of the concatenated fractions, with the remaining 95 % used for phosphopeptide enrichment using immobilized metal affinity chromatography (IMAC). Further detail on the phosphopeptide enrichment process is provided here[5]. Flow through from the IMAC was further processed for acetylpeptide enrichment using an acetyl-lysine antibody (Cell Signaling Technologies, #13416). Four to 5 µl of each fraction were used for LC-MS/MS analysis for each proteomic assay.

The LC-MS/MS analysis was conducted in somewhat different ways for heart and liver, compared to gastrocnemius, white adipose, lung, kidney and cortex due to site-specific differences. For heart and liver, the online separation was performed using a nanoflow Proxeon EASY-nLC 1200 UHPLC system (Thermo Fisher Scientific) and an in-house packed 22 cm x 75 µm C18 silica picofrit capillary column used for the LC step. For global proteome, 1 µg was loaded in a 2µL volume, whereas for phosphoproteome, acetylome, and ubiquitylome, a 4-µl volume was used with 50% of each fraction sample. Mass-spectrometry analysis was conducted using a Q-Exactive Plus mass spectrometer (Thermo Fisher Scientific) for the global proteome, and a Q-Exactive HFX mass spectrometer (Thermo Fisher Scientific) for the phosphoproteome, acetylome and ubiquitylome. Specific analysis settings have been described elsewhere[5]. For the other tissues, analyzed at the Pacific Northwest National Laboratory, online separation was instead performed using a nanoAcquity M-Class UHPLC system (Waters), and an in-house packed, 25 cm x 75 µm C18 silica picofrit column used for the LC step. The mass-spectrometry analysis was conducted using a Q Exactive HF mass spectrometer (Thermo Fisher Scientific) for the global proteome. For the phosphoproteome, a Dionex Ultimate 3000 UHPLC direct-inject system (Thermo) was used for online separation, a 30 cm x 75 µm C18 silica picofrit column for the LC and Q-Exactive HFX mass spectrometer (Thermo Fisher Scientific) for the mass spectrometry analysis. Additional details on the sample processing is available elsewhere[5]. Acetylome and ubiquitylome was only analyzed in the heart and the liver.

Quantification of all proteomic analysis were calculated as $\log_2$ TMT ratios to the common reference, with analytes not fully quantified in at least two plexes within a tissue removed. Sample normalization was performed by median-centering and mean absolute deviation scaling. Limma in R (v 3.48.0) was used to remove batch effects. Correction for protein abundance in ubiquitylome was accomplished by fitting a global linear model between a site-specific PTM and the cognate protein and extracting the residuals.

#### Non-targeted metabolomics

Non-targeted metabolomics was performed through hydrophilic interaction liquid chromatography (HILIC) analyses at the Broad Institute of MIT and Harvard, and through reverse-phase and ion pairing profiling at the University of Michigan. For HILIC positive analyses, 10 mg of powdered tissue was homogenized in 300 µL of 10/67.4/22.4/0.018 v/v/v/v water/acetonitrile/methanol/formic acid containing stable isotope-labeled internal standards. For plasma, 10 µl mixed with 90 µL of 74.9/24.9/0.2 v/v/v acetonitrile/methanol/formic acid, was used. After centrifugation for 10 minutes at 9,000 x g, 4°C, samples were injected onto a HILIC column and eluted at 250 µL/min with mobile phase A for 30 seconds, and subsequently with a linear gradient to 40 % mobile phase B for 10 minutes, with stable elution for 4.5 minutes after that. MS analyses using electrospray ionization was conducted in the positive ion mode using full scan analysis over 70-800 *m/z* at 70,000 resolution and 3 Hz data acquisition rate using a Q-Exactive hybrid quadrupole Orbitrap mass spectrometer (Thermo Fisher Scientific). TraceFinder software (Thermo Fisher Scientific) was used to process raw data for targeted peak integration and Progenesis QI (Nonlinear Dynamics, Waters) for peak detection and integration of metabolites of known and unknown identity.

For reverse phase and ion-pairing analyses, 50 µl plasma was mixed with 200 µL of extraction solvent (1:1:1 v:v methanol:acetonitrile:acetone containing internal standards and homogenized through vortexing for 10 seconds. Details about internal standard concentrations are provided elsewhere[5]. Non-powdered solid tissues were weighed and mixed with 1:1:1:1 methanol:acetonitrile:acetone:water in a ratio of 1 ml per 50 mg tissue and homogenized using a sonicator. Subsequent steps were the same for plasma and tissues and included a 10 min incubation on ice, followed by centrifugation at 15 000 x g. The supernatant (300 µL for tissues and 150 µL for plasma) was dried with a nitrogen blower and reconstituted in water:methanol (8:2 v:v) for the LC-MS analysis. The reverse phase analyses were conducted on an Agilent 1290 Infinity II / 6545 qTOF MS system with a JetStream electrospray ionization source (Agilent Technologies) using a Waters Acquity HSS T3 column (Waters Corporation). Injection volume was 5 µL and flow rate 0.45 ml/min, mobile phase A water with 0.1% formic acid, and mobile phase B methanol with 0,025% formic acid. Each sample was analyzed in both the positive and negative ion mode. The ion pairing analyses were conducted using an Agilent Zorbax Extend C18 1.8 µm RRHD column, 2.1 x 150 mm ID, equipped with a matched guard column. Mobile phase A consisted of 97% water and 3% methanol, mobile phase B 100% methanol and mobile phase C 100% acetonitrile. Phase A and B also contained 15 mM tributylamine and 10 mM acetic acid. Injection volume was 5 µL, with variable flow rate and time (for details, see [5]). Mass spectrometry analysis was conducted in the negative ion mode. Profinder v8.0 (Agilent Technologies) was used for targeted compound detection and relative quantitation, while custom scripts were used for non-targeted feature detection. Agilent Mass Profiler Pro (v8.0) and Masshunter Qualitative Analysis were used for feature alignment and recursive feature detection. Prior to normalization, features missing from >50% of all samples in a batch or >30% of QC samples were removed. Data reduction was then performed using Binner [6] and normalized using the Systematic Error Removal Using Random Forest approach [7]. The performance of the normalization was validated using relative SD for QC samples.

#### Non-targeted lipidomics

Non-targeted lipidomics was performed using 10 mg of powdered tissue or 25 µL of plasma. Tissue was homogenized in 400 µl isopropanol (containing stable isotope-labeled internal standards from Avanti Polar Lipids (Alabaster)), using freeze thawing in liquid nitrogen and sonication, while plasma was only mixed with 75 µL isopropanol and internal standards. Samples were then centrifuged for 5 min at 21,100 x g and supernatants used for LC-MS on a Vanquish chromatograph with an Accucore C30 column (2.1 × 150 mm, 2.6 µm particle size), coupled to a high-resolution accurate mass Q-Exactive HF Orbitrap mass spectrometer (all from ThermoFisher Scientific). Injection volume was 2 µL, mobile phase A was 40:60 water:acetonitrile with 10 mM ammonium formate and 0.1% formic acid, and mobile phase B 10:90 acetonitrile:isopropyl alcohol, with 10 mM ammonium formate and 0.1% formic acid. Details on the gradient program are provided elsewhere[5]. Full mass spectrometry data was acquired with 240,000 resolution over the 150-2000 m/z range. Raw LC-MS data was processed using Compound Discoverer V3.0 (ThermoFisher Scientific). Peak area was corrected for QC sample peak areas across the batch, followed by filtering with background and QC filters. Features absent in >50% of the QC pooled injections and a coefficient of variance

<30 % were removed. Annotations were based on mass and relative abundance, retention time and MS2 pattern.

### Targeted metabolomics and lipidomics

Targeted assays were performed for branched-chain keto acids, acyl CoA's and nucleotides at Duke University, and for amino acids and amino metabolites, TCA cycle metabolites, ceramides and acylcarnitines at the Mayo Clinic. Targeted lipidomics was performed at Emory University. Ten µl plasma and 200 µl tissue homogenate was used for analysis of branched-chain keto acids, which were extracted using ethyl acetate as described previously [8]. Five hundred µl tissue homogenate was used for acyl CoA extraction, as reported previously for liquid phase [9] and solid phase [10] extraction. Nucleotides were extracted as previously described [11,12]. Samples were subsequently centrifuged at 14,000 x g for 5 minutes and supernatant used for LC–MS/MS. Extracts for branched-chain keto acids, acyl CoA's and nucleotides were analyzed on a Xevo TQ-S triple quadrupole mass spectrometer (Waters) and endogenous levels were quantified using calibrators by spiking tissue homogenates (Acyl CoA's and nucleotides) or fetal bovine serum (keto acids) with authentic analytes (all from Sigma-Aldrich).

For targeted amino acid and amino metabolite profiling, 20 ml of plasma or 5 mg of tissue homogenate was utilized. Extraction and analysis was conducted as previously described [13,14]. Targeted profiling of ceramides and sphingolipids, using 25 µl plasma or 5 mg of tissue homogenate, has also been previously reported [15,16], as have the targeted profiling of acylcarnitines[17,18] (specifically C0-C18:1), which was also performed using LC-MS/MS from the same amount of plasma and tissue. TCA metabolites were profiled from 5 mg of tissue or 50 µl plasma using gas chromatography mass-spectrometry, as previously described [18,19] with minor modifications reported elsewhere [5].

Targeted lipidomics was performed on 10mg of powdered tissue, homogenized in 100 µl PBS, followed by dilution with 100 µL 20 % methanol and spiked with 1 % BHT solution according to previously used methods[20,21]. After centrifugation at 14,000 rpm for 10 min, the supernatant was transferred to 96-well plates and loaded onto C18 SPE columns and eluted with 400 µl methyl formate. External standards were all purchased from Cayman Chemical. An ExionLC (SCIEX) chromatograph with an AccucoreTM C18 column (ThermoFisher), and a SCIEX QTRAP 5500 mass spectrometer was used for LC-MS/MS. Mobile phase A was water with 10mM ammonium acetate, and mobile phase B acetonitrile with 10 mM ammonium acetate. Details on the gradient program and subsequent mass spectrometry is provided elsewhere [5]. Raw data was processed using Sciex OS (AB SCIEX, Version 1.6.1).

### Metabolomics and lipidomics data processing and normalization

All metabolomics datasets were partitioned into named compounds for analytes that were confidently identified, and unnamed for those without a standard chemical name. Only named metabolites were included in this analysis, which included $\log_2$-transforming the data and removing analytes with >20% missing values (including negative values). For targeted datasets with > 12 analytes and all untargeted datasets, remaining missing values were imputed using K-nearest neighbors (k=10 samples). Outlier identification was done using the sample-level median correlation against the other samples, and all 21 confirmed technical outliers were manually reviewed by the metabolomics sites. Only untargeted datasets were normalized, using median-centering. Metabolites that were identified by two or more platforms (1116 metabolites in total) were integrated using meta-regression.
